# Integrative In Silico and Experimental Identification of Non-Covalent UBE2N Inhibitors Enhancing PARP Inhibitor Sensitivity

**DOI:** 10.64898/2025.12.09.693205

**Authors:** Côme Ghadi, Shafi Ullah Khan, Léonie Ibazizene, Florian Schwalen, Charline Kieffer, Peggy Suzanne, Julie Jaouen, Hassiba Bouafia, Xavier Thuru, Matthieu Meryet-Figuiere, Louis-Bastien Weiswald, Anne Sophie Voisin-Chiret, Jana Sopkova-de Oliveira Santos

## Abstract

UBE2N, an E2 ubiquitin-conjugating enzyme, has emerged as a compelling therapeutic target in oncology due to its critical roles in DNA damage repair and NF-κB signalling. While covalent inhibitors have shown preclinical promise, non-covalent inhibitors offer potential advantages in terms of selectivity and reduced off-target effects. However, structural and mechanistic data for non-covalent UBE2N inhibitors remain scarce. To address this gap, we implemented a dual *in silico* strategy combining structure-based molecular docking and ligand-based 3D pharmacophore modelling. Screening a home library of ∼19,000 compounds targeting both the ubiquitin-binding and cofactor interfaces of UBE2N, we identified 22 candidates suitable for biological evaluation.

Among these, two compounds, CERMN-2 and CERMN-16, emerged as promising non-covalent inhibitors. CERMN-16, structurally related to the natural compound Variabine B (identified through 3D pharmacophore screening), significantly reduced SKOV-3 ovarian cancer cell viability and enhanced their sensitivity to the PARP inhibitor Olaparib. CERMN-2, identified through docking, also demonstrated a synergistic effect with Olaparib and showed low toxicity in normal ovarian epithelial cells. Molecular dynamics simulations indicated distinct binding modes for each compound, consistent with their targeted binding sites. Biophysical experiments revealed weak binding of CERMN-16 to UBE2N, whereas CERMN-2 bound UBE2N in two orthogonal assays (Microscale thermophoresis and Nano differential scanning fluorimetry). CERMN-16, and more importantly CERMN-2, therefore represent promising leads for the development of selective, non-natural, non-covalent UBE2N inhibitors. These results provide new insights into UBE2N inhibition and support further investigation of their mechanisms of action and therapeutic potential in combination cancer therapies.

**HIGHLIGHTS:**

- Dual *in silico* screening (docking and 3D pharmacophore) identified new non-covalent UBE2N inhibitor candidates.
- Two compounds, CERMN-2 and CERMN-16, displayed synergistic activity with Olaparib in ovarian cancer cells.
- MD simulations revealed distinct, site-specific binding modes for both compounds.
- Biophysical assays confirmed UBE2N binding for CERMN-2, identifying it as a promising non-natural, non-covalent lead.

## INTRODUCTION

According to the most recent estimates, cancer accounts for 20 million cases and 9.7 million deaths worldwide annually.^1^ Despite continuous advances in treatments, the persistence of high rates of death in cancer patients underlines the need for new therapeutic options. In most cases, therapeutic strategies involve surgical resection (for solid cancers) associated with DNA-damaging chemotherapy or radiotherapy. One of the major challenges in cancer therapy is resistance to treatments, whether intrinsic or acquired during the course of therapy.^2^

Sensitizing tumours to DNA damaging therapeutics could therefore improve patient outcome. DNA double-strand breaks, the most lethal form of DNA damage, are primarily repaired by two pathways: Non-Homologous End Joining (NHEJ) and Homologous Recombination (HR). While HR is considered faithful, NHEJ is more error-prone and often results in large genomic rearrangements and cell death. Hence, defects in the HR pathway increase sensitivity to DNA-damaging drugs, as evidenced by the enhanced response to carboplatin and PARP inhibitors (PARPi) in ovarian cancer patients with impaired HR. This HR deficiency (HRD) phenotype can arise from *BRCA1/2* mutations or from alterations in other genes within this complex pathway.^3^

Several strategies are currently being explored to impair HR pathway, such as inhibition of LSD1 (Lysine-specific histone demethylase 1A), which reduces *BRCA1/2* transcription^4^ or saturation of the DNA damage response pathways.^5^ Other approaches include inhibiting the ATR kinase, which acts early in the DNA damage response^6^ or targeting HSP90.^7^ These latter two strategies are currently in clinical trial phases, although none have yet received regulatory approval.^7,8^

These approaches target various steps of the complex machinery of HR DNA repair. We chose to focus our attention on UBE2N, a key factor involved in both HR and post-replicative DNA repair, two pathways critical for cellular response to various DNA damaging agents, such as PARPi or carboplatin.^9^ This dual involvement mirrors the functions of BRCA1, involved as well in HR and post-replicative repair.^10^ In this regard, inhibiting UBE2N might closely mimic the HRD phenotype. In the nucleus, its functions in DNA repair depend on interaction with its cofactor, UBE2V2. Meanwhile, in the cytoplasm, UBE2N positively regulates NF-κB signaling, a process that requires its association with the cofactor UBE2V1.^9^

UBE2N inhibition has been investigated across several malignancies, including melanoma,^11^ lymphoma,^12^ prostate cancer,^13^ and others, primarily due to its role in the NF-κB pathway. However, to our knowledge, its association with DNA-targeting drugs has been reported only once. In a study we conducted, UBE2N inhibition was shown to sensitize several clinically relevant Patient-Derived Tumour Organoid models of ovarian cancer to carboplatin.^14^

UBE2N is an E2 ubiquitin-conjugating enzyme, at play in the ubiquitination process. This process involves tagging target proteins with chains of ubiquitin, a small regulatory protein, to alter their function, localization, or stability. The process begins with the ubiquitin-activating enzyme (E1), which activates ubiquitin in an ATP-dependent manner, forming a high-energy thioester bond between ubiquitin and the E1 enzyme. The activated ubiquitin is subsequently transferred from the E1 enzyme to the active site cysteine (Cys87) of UBE2N via a thioester bond. UBE2N operates in collaboration with its cofactors - UBE2V1 or UBE2V2 - and works alongside E3 ubiquitin ligases to catalyse the transfer of K63-linked ubiquitin chains to target proteins.^9^

The catalytic ubiquitinating activity of UBE2N is naturally inhibited by the deubiquitinating enzyme OTUB1 (OTU domain-containing ubiquitin aldehyde-binding protein 1). UBE2N forms a non-catalytic, inhibitory complex with OTUB1 that blocks ubiquitin transfer.^15^ In addition to this natural regulation, UBE2N catalytic activity can also be inhibited by synthetic compounds.

To date, only a few UBE2N inhibitors (UBE2Ni) have been identified, with covalent inhibitors being the most extensively studied. Among these, NSC697923 is a small-molecule inhibitor that covalently binds to the Cys87 residue in the active site of UBE2N via a Michael addition, attaching a 5-nitrofuran moiety. This compound effectively inhibits UBE2N activity, blocking DNA damage responses and NF-κB signaling.^12^ Similarly, BAY 11-7082 targets Cys87 in UBE2N via the same mechanism, forming a smaller chemical adduct. However, its broad reactivity toward sulfhydryl groups raises concerns about off-target effects and toxicity.^16^ UC-764864 and UC-764865, two structurally related compounds, also covalently modify Cys87 in UBE2N and suppress UBE2N-dependent signalling in leukaemia cells. Despite their promising activity, data on their selectivity and potential off-target effects remain limited, highlighting the need for further investigation.^17^

In addition to covalent inhibitors, non-covalent compounds represent a valuable alternative strategy for targeting UBE2N, offering the potential for improved selectivity, although sometimes at the expense of potency. Among these, Leucettamol A has been reported to disrupt the UBE2N/UBE2V1 interaction; however, its use is limited by chemical instability and isomerization.^18^ Similarly, Manadosterols A and B have been shown to inhibit the UBE2N/UBE2V1 interaction.^19^ Variabine B, a β-carboline alkaloid, interferes with the interaction between UBE2N and its cofactors UBE2V1 or UBE2V2, exhibiting an IC₅₀ of 4 µg/mL (approximately 16 µM).^20^ ML307 has emerged as a potent submicromolar inhibitor of UBE2N enzymatic activity; however, its poor microsomal stability renders it unsuitable for *in vivo* application.^21^ In parallel, peptoid scaffolds are being explored as a novel class of non-covalent UBE2N inhibitors, representing a promising direction for therapeutic development.^22^

While significant work has been made in identifying UBE2Ni, most of them have been investigated in a few studies, or even a single one, with the exception of NSC697923. Moreover, none of the existing compounds appear to exhibit the necessary properties for potential clinical application in humans. Consequently, major challenges persist, and further research is required to develop highly selective and effective UBE2N inhibitors suitable for therapeutic use. Given that the non-covalent binding mode of UBE2Ni has not been fully explored, we developed a robust *in silico* approach combining docking and 3D pharmacophore modelling, further refined through molecular dynamics simulations. This model enabled systematic virtual screening of our home chemical library (CERMN chemolibrary), which comprises approximately 19,000 compounds, leading to the identification of several promising candidates.

## MATERIALS AND METHODS

### Virtual screening

#### 3D structures and PCA analysis

All the available 3D structures of UBE2N were retrieved from the PDB database^23^ and visually analysed using PyMOL software. PCA analyses were conducted using a custom R script with the bio3d library.^24^ For the PCA analysis, we selected X-ray structures without mutations and without gaps in their sequence from the available dataset, corresponding to twenty structures (PDB IDs: 1J7D, 3HCT, 3HCU, 3VON, 3W31, 4DHI, 4DHJ, 4DHZ, 4IP3, 4ONM, 4ONN, 4ORH, 5YWR, 6KFP, 6KG6, 6LP2, 6P5B, 6S53, 6UMP, and 7BXG). The analysis was performed on the common sequence from residue 5 to residue 150, focusing on the Cα atoms to annotate the skeletal motion of the protein.

#### ML307 and Variabine B docking and CERMN chemolibrary screening by docking

The blind docking of ML307 and Variabine B was performed using AutoDockVina software (v1.1.2)^25^ on three structures representing different shapes of the ubiquitin binding site: 3HCU,^26^ 4ONM,^16^ and 6UMP.^27^ To prepare the structures for docking, ions, water molecules, and co-crystallized ligands and partners were removed, and one UBE2N monomer was retained. The histidine residues were modelled according to PropKa 3.0 software prediction at physiological pH=7.4: His36 as HSE and His77 as HSD.^28,29^ Polar hydrogen atoms were added using AutoDockTools on the protein, and partial charges were calculated with the Gasteiger algorithm.^30^

The 3D models for ML307 and Variabine B were built using Marvin Sketch from the ChemAxon Package (http://www.chemaxon.com). The preferred protonation state at pH 7.4 was predicted by MarvinView from the same package, and the predicted major form was used in the docking studies. During the blind docking procedure, the default exhaustiveness value of 8 was applied and 20 different poses were generated for each ligand. The box parameters used in the blind docking are summarized in Supplementary Information (SI) Table S1.

To screen the CERMN chemolibrary, we began by preparing the library’s compounds. Salts were removed, and the compounds were converted to 3D structures. Protonation states at pH 7.4 were predicted using Open Babel 3.0.1,^31^ which was also employed to convert the compound coordinates into PDBQT format. The chemolibrary was then screened with AutoDock Vina with an exhaustiveness value set to 8.^25^ The box parameters for the screening process are detailed in Table S1. For each ligand, 10 distinct poses were generated. The highest-scoring pose of each ligand was used as a criterion for their selection for further analysis. For the selected ligand, all 10 poses were visually inspected to confirm the ligand binding to the target site and to assess the plausibility of their binding modes.

#### 3D pharmacophore screening

The 3D pharmacophores were generated using LigandScout 4.4.9 software^32,33^ through two approaches. The first was a ligand-based approach, utilizing Variabine B, a natural non-covalent inhibitor of the UBE2N/UBE2V2 interaction, derived from the marine sponge *Luffariella variabilis*.^20^ The pharmacophore generated from Variabine B was then used to screen the CERMN chemolibrary.

In the second approach, we used LigandScout to generate 3D pharmacophore models directly from the topology of the binding sites. First, the “Calculate Pocket” function was applied to identify druggable pockets on the protein surface of the three crystallographic structures selected previously for docking screening. The default threshold parameter of 0.3 was used for pocket detection across all cases, while the Buriedness parameter ranged from 0.2 to 0, depending on the structure (see SI Table S2). Next, the “Apo Site Grid” function was applied to the selected interaction sites, either with ubiquitin or UBE2V2/UBE2V1, to compute all potential interactions within the pocket. Finally, the “Apo Site Pharmacophore” tool, with default settings (Buriedness at 0.7 and surface grid at 0.19), was employed to create 3D pharmacophores based on the previously identified interactions. The pharmacophore features situated far from the target sites were removed.

To perform the 3D pharmacophore-based virtual screening, the processed CERMN library was imported into LigandScout. Conformers were generated using the iCon conformer generator implemented in LigandScout, with the ’BEST’ option selected to ensure high-quality conformer generation.^34^ Pharmacophore screening was conducted using the Pharmacophore-Fit scoring function. The “Match All Query Features” option was selected, and the retrieval mode was set to stop after the first matching conformation. The maximum number of omitted features varied depending on the pharmacophore model. Exclusion volume checks were enabled and validated during the screening process.

#### Molecular Dynamics

All dynamics simulations were performed using NAMD 2.13.^35^ The all-atom CHARMM36m forcefield^36,37^ was used for the UBE2N protein and CGENFF^38^ one for the selected ligands. Moreover, the additional force field of hydrogen mass repartition (HMR)^39^ was applied in each simulation. The starting systems were generated by the CHARMM-GUI server^40,41^. The histidine residues in UBE2N were modelled in the same state as for docking. The *N* and *C*-termini of protein domains were capped. Each system was solvated using the TIP3P explicit water model^42^ within a rectangular box; the box size ensured that the simulated complex was at a minimum distance of 10 Å from the edge. To neutralize the total charge system, 0.15 M of KCl was added. The vacuum dielectric constant was used during all calculations. Cubic periodic boundary conditions were applied to the systems by using the IMAGE algorithm. The applied cut-off distance was 16 Å, and van der Waals interactions were truncated using a force-switching function between 10 and 12 Å. The Particle Mesh Ewald (PME) was used to calculate long-range electrostatic interactions.^43^ The SHAKE algorithm was applied to restrain all bonds involving hydrogen atoms.

Firstly, each simulation underwent energy minimization in 10,000 steps, with harmonic restraints applied on heavy atoms (1 kcal/(mol×Å^2^) force constant for backbone atoms and 0.5 kcal/(mol×Å^2^) force constant for sidechain atoms). Next, the minimized systems were heated to 303.15 K, and the dynamics were temperature-equilibrated during 250 ps *via* heating reassignment under NVT conditions using a time step of 2 fs, with harmonic restraints applied to heavy atoms. Finally, the systems ran freely for 100 ns under NPT conditions, with a time step of 4 fs. Langevin dynamics with a damping coefficient of 1 ps^−1^ was used to maintain the system temperature, and the Nosé–Hover–Langevin piston method to control the pressure at 1 atm. Production trajectories were saved every 100 ps, and subsequent analyses were performed using the CHARMM c40b2 homemade scripts.^44^ The MD simulations were performed in triplicate for each compound and each pose (ML307, Variabine B, CERMN-2), except for those used to validate the compounds after screening.

### Biological evaluation

#### Cell proliferation and cell viability of selected compounds from CERMN chemolibrary

SKOV-3, a human ovarian carcinoma cell line, was purchased from the American Type Culture Collection (ATCC, Manassas, VA, USA). Cells were maintained in RPMI 1640 medium (Thermo Fisher Scientific, Waltham, MA, USA) supplemented with 2 mM GlutamaxTM (Thermo Fisher Scientific), 25 mM HEPES (Thermo Fisher Scientific), 10% fetal bovine serum (FBS; HyClone, GE Healthcare Life Sciences, Logan, UT, USA), and 33 mM sodium bicarbonate (Sigma-Aldrich, St. Louis, MO, USA). Epithelial T1074 cells were obtained from Applied Biological Materials (Abm, Vancouver, Canada) and supplemented with 2 mM Glutamax, 25 µM HEPES, 10% fetal calf serum and 33 mM sodium bicarbonate. Cultures were maintained at 37 °C in a humidified incubator with 5% CO_2_. Cells were passaged at approximately 80% confluence using 0.25% trypsin-EDTA (Thermo Fisher Scientific). The UBE2N inhibitor (NSC697923, Cat. No.: HY-13811) and the PARP inhibitor (Olaparib, Cat. No.: HY-10162) were purchased from MedChemExpress. ML307 (3R)-N-({1-[(3chlorophenyl)methyl]piperidin4-yl}methyl)-1-{6H,7H,8H,9Hpyrido[2,1-h]purin-4yl}piperidine-3-carboxamide (Cat. No.: Z8994743401) was custom-synthesized by Enamine, while Variabine-B was synthesized in-house (see SI). Library and stock solutions were prepared in sterile dimethyl sulfoxide (DMSO; Sigma-Aldrich) according to the manufacturers’ instructions, reaching a final concentration of 50 mM, and stored at -80 °C.

#### Treatment Protocol

SKOV-3 in and T1074 cells were seeded at a density of 1 × 10^3^ cells per well in 96-well plates and allowed to adhere for 24 hours. Subsequently, cells were exposed to increasing concentrations (5, 10, 20 and 40 µM) of the test compounds, either alone or in combination with Olaparib (10 µM), for 72 hours.

#### Cell Viability Assay

Cell metabolic activity was used as a proxy for cell viability the CellTiter 96® Aqueous MTS Reagent Powder (Promega, Madison, WI, USA) according to the manufacturer’s instructions. Briefly, after treatment duration of 72 hours, 20 µL of freshly prepared MTS reagent (2 mg/mL MTS and 0.92 mg/mL PMS in DPBS) was added to each well, and the plates were incubated for further 2 hours in dark. The absorbance was measured at a 490 nm wavelength using a microplate reader (FLUOstar OPTIMA, BMG). Cell viability was directly proportional to the measured absorbance.

#### Colony forming assay

SKOV3 cells were seeded at 1 × 10³ cells per well in 6-well plates containing 2 mL of RPMI medium. Following overnight incubation at 37 °C with 5% CO₂ to allow cell attachment, the medium was replaced with fresh medium containing the designated treatments at predetermined concentrations. The plates were incubated for 7 days at 37 °C with 5% CO₂. Colonies were stained with 0.25% Crystal Violet solution in Methanol for 20 minutes. Colonies were counted manually.

Data analysis was performed using GraphPad Prism 9 software (GraphPad Software Inc.) Results from all experiments were reported as the means ± standard deviation obtained from three independent experiments. A one-way ANOVA was employed for comparisons.

#### Nano differential scanning fluorimetry (nanoDSF)

NanoDSF measurements were performed using a Tycho NT.6 instrument (NanoTemper Technologies GmbH). His-tagged UBE2N (Novus Biologicals) was diluted to 3 µM in assay buffer (PBS pH 7.4 with 0,05% Tween20) and centrifugated for 10 min at 20 000 g and at 4°C. The protein solution was then mixed with the ligand (1:1, v/v) at the desired concentration, resulting in a final DMSO concentration of 2%. The mixture was incubated for 20 min at room temperature. Mixture was then transferred into capillaries. Thermal protein denaturation was monitored by measuring fluorescence intensity ratio at 350/330 nm under a temperature ramp from 35 to 95 °C (30 °C/min). Melting temperatures (Tm) values were determined using the internal Tycho NT.6 analysis software. Each condition was measured in triplicate.

#### Microscale thermophoresis (MST)

Microscale thermophoresis (MST) experiments were performed using a Monolith NT.115 Pico instrument (NanoTemper Technologies GmbH) equipped with red and blue filter sets. His-tagged UBE2N (Novus Biologicals) was diluted to 200 nM in assay buffer (PBS pH 7.4 with 0.05% Tween20). The protein was labelled using the His-Tag Labeling Kit RED-tris-NTA 2nd generation (NanoTemper Technologies), at a final concentration of 100 nM, according to the manufacturer’s instructions.

Ligands were serially diluted (1:1) over 16 gradient steps. The labelled protein (10 nM) was added to each ligand dilution at a 1:1 ratio and incubated for 15 min at room temperature. Standard capillaries (NanoTemper) were individually filled and loaded into the instrument. Data were acquired using high MST power and 20% LED with “MO.Control” software (NanoTemper). Measurements were analysed using “MO.Affinity Analysis” software (NanoTemper).

## RESULTS

### Analysis of UBE2N 3D structures

Numerous crystal structures of the UBE2N protein in complex with various partners have been determined using X-ray diffraction. To date, 40 UBE2N-containing structures have been deposited in the Protein Data Bank (PDB).^45^ Analysis of these structures - including complexes with UBE2V2, UBE2V1, ubiquitin, E3 ligases, tumour-associated factors, and bacterial proteins - has enabled the identification of key interaction interfaces on UBE2N (Figure 1).

**Figure 1.**
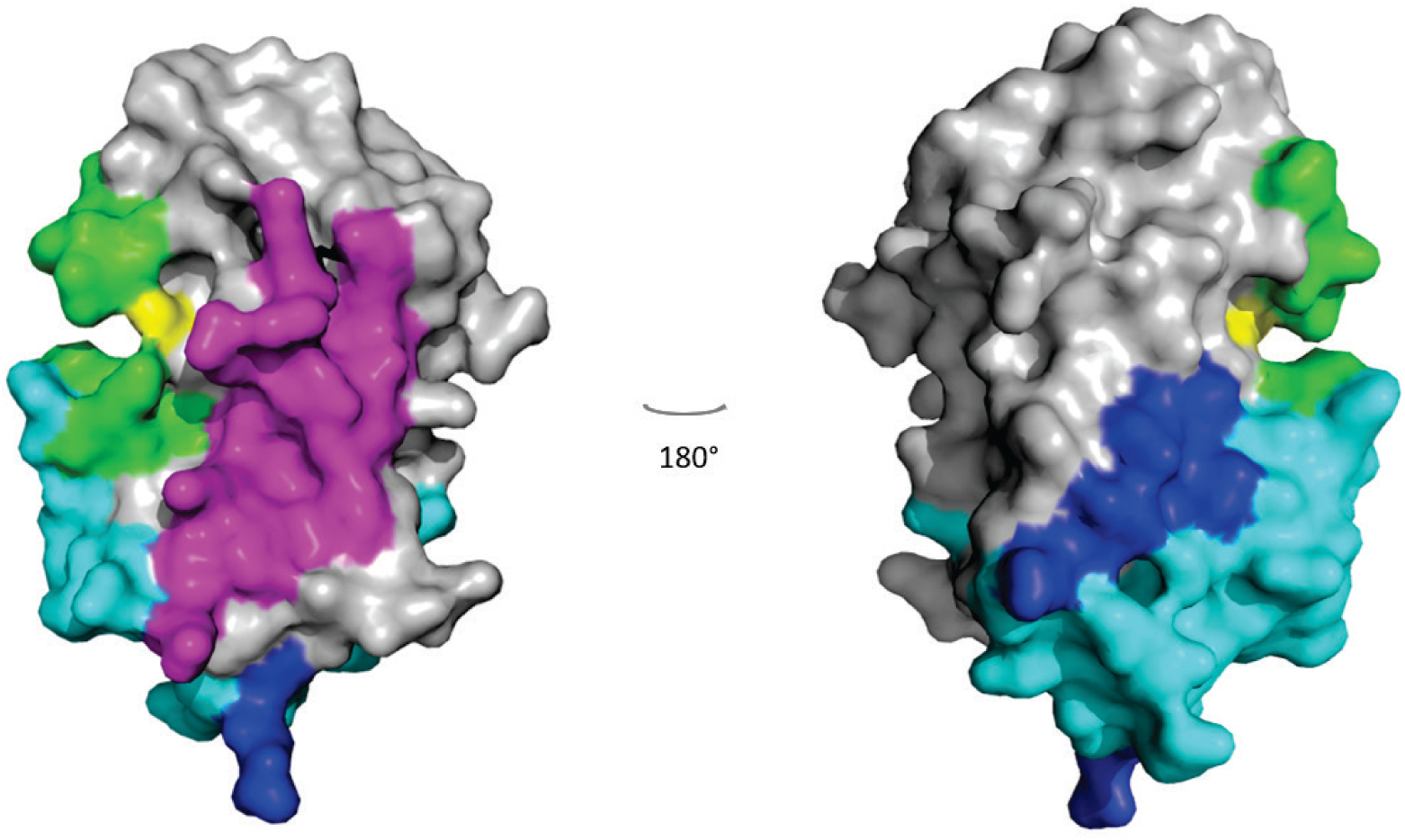
3D Structure of UBE2N and Its Interactions with Various Partners. Two views of surface representations of UBE2N showing interaction zones with its various partners: magenta - regions interacting with UBE2V2 and UBE2V1; green - regions interacting with Ubiquitin; cyan - regions interacting with E3 ligases and tumour-related factors; blue - regions interacting with bacterial proteins; yellow - Cys87 residue. The deubiquitinating enzyme OTUB1 was found bound either in the cyan or magenta regions in the solved complex structures.

The PDB structures complexed with ubiquitin reveal that the UBE2N active site (shown in green in Figure 1) is a cavity near Cys87 (yellow in Figure 1). To catalyse the elongation of K63-type Ub chains, UBE2N requires a partner protein, either UBE2V1 or UBE2V2, with which it forms heterodimers. The interaction surface with UBE2V2 or UBE2V1 is distinct from that of ubiquitin (coloured in pink in Figure 1), and it is identical for both E2 ligases, as the residues of UBE2V2 and UBE2V1 that interact with UBE2N are strictly conserved.^22^ Therefore, UBE2N inhibitors that affect the interaction with either of these partners would likely impact both, which would be advantageous given the functional importance of these interactions.

To date, no UBE2N structure co-crystallized with a non-covalent synthetic inhibitor has been solved. There are, however, four structures co-crystallized with the deubiquitinating enzyme OTUB1,^46,15^ and two structures of UBE2N co-crystallized with covalent inhibitors: NSC697923 (PDB ID: 4ONM) and BAY 11-7082 (PDB ID: 4ONN)^16^. The X-ray structures reveal that OTUB1 can bind either at the UBE2V2/UBE2V1 binding site or the E3 ligase binding site (Figure 1), making its mode of action unclear.

The interest in the X-ray structures of covalent inhibitor/UBE2N complexes is limited. For instance, NSC697923 binds to UBE2N through a covalent bond to Cys87, and only the 5-nitrofuran moiety added to the sulfhydryl group of Cys87 is visible in the crystal structure.^16^ Despite this, the solved structure showed a conformational change in the loop between amino acids 114-124 in UBE2N, creating a cavity adjacent to Cys87. The NSC697923 nitrofuran moiety present in the crystal lodges in this cavity and establishes a hydrogen bond with Asn123. Similar observations were made for BAY 11-7082, the other covalent UBE2N inhibitor. In this structure, the prop-2-enenitrile moiety is added to the Cys87 sulphur atom and lodges in the same cavity.

These structures emphasize that the shape of UBE2N’s active site near Cys87 is influenced by the movement of the flexible 114-124 loop, which modulates the cavity volume. This modulation may play a key role in ubiquitin inhibitors selectivity for UBE2N.

### Docking screening of CERMN chemolibrary

Since the motion of the 114-124 loop affects the ubiquitin-binding cavity - observed in structures co-crystallized with covalent inhibitors and hypothesized to contribute to selectivity-we first applied Principal Component Analysis (PCA) to available UBE2N X-ray structures. This analysis enabled us to identify different conformations of the 114-124 loop and to assess cavity variability among all available X-ray structures of UBE2N. PCA extracts principal components (PCs) that capture the highest variance within the dataset, highlighting key regions of structural flexibility in UBE2N. The first component (PC1) captured the transformation of the α-helix spanning residues 89-92 into a flexible and disordered loop, located relatively far from the ubiquitin-binding site (Supporting information Figure S1). The second component (PC2) primarily reflected the motion of the loop between residues 114-124 (Figure S1A).

To capture the full range of loop conformations and the shape of the ubiquitin-binding site, we selected three structures for our modelling studies: one with a high positive PC2 value (3HCU), one with a value close to zero (4ONM), and one with a significantly negative value (6UMP) (Figure 2A). We selected the 3HCU structure over 3VON because the mobile segment of the loop corresponds to the same residues as in 4ONM and 6UMP (Figure S1B). Specifically, 3HCU represents a complex of UBE2N with TRAF2^26^, 4ONM corresponds to a UBE2N/UBE2V2 structure co-crystallized with the covalent inhibitor NSC697923^16^, and 6UMP is a UBE2N/ubiquitin complex bound to the *Legionella pneumophila* bacterial protein MavC^27^. Since the ubiquitin-binding site and the UBE2V2/UBE2V1 interaction site are adjacent on the UBE2N surface, structural changes in the ubiquitin-binding site also influence the UBE2V2/UBE2V1 site. Consequently, all docking studies targeting both sites were conducted independently using the three selected UBE2N structures.

**Figure 2:**
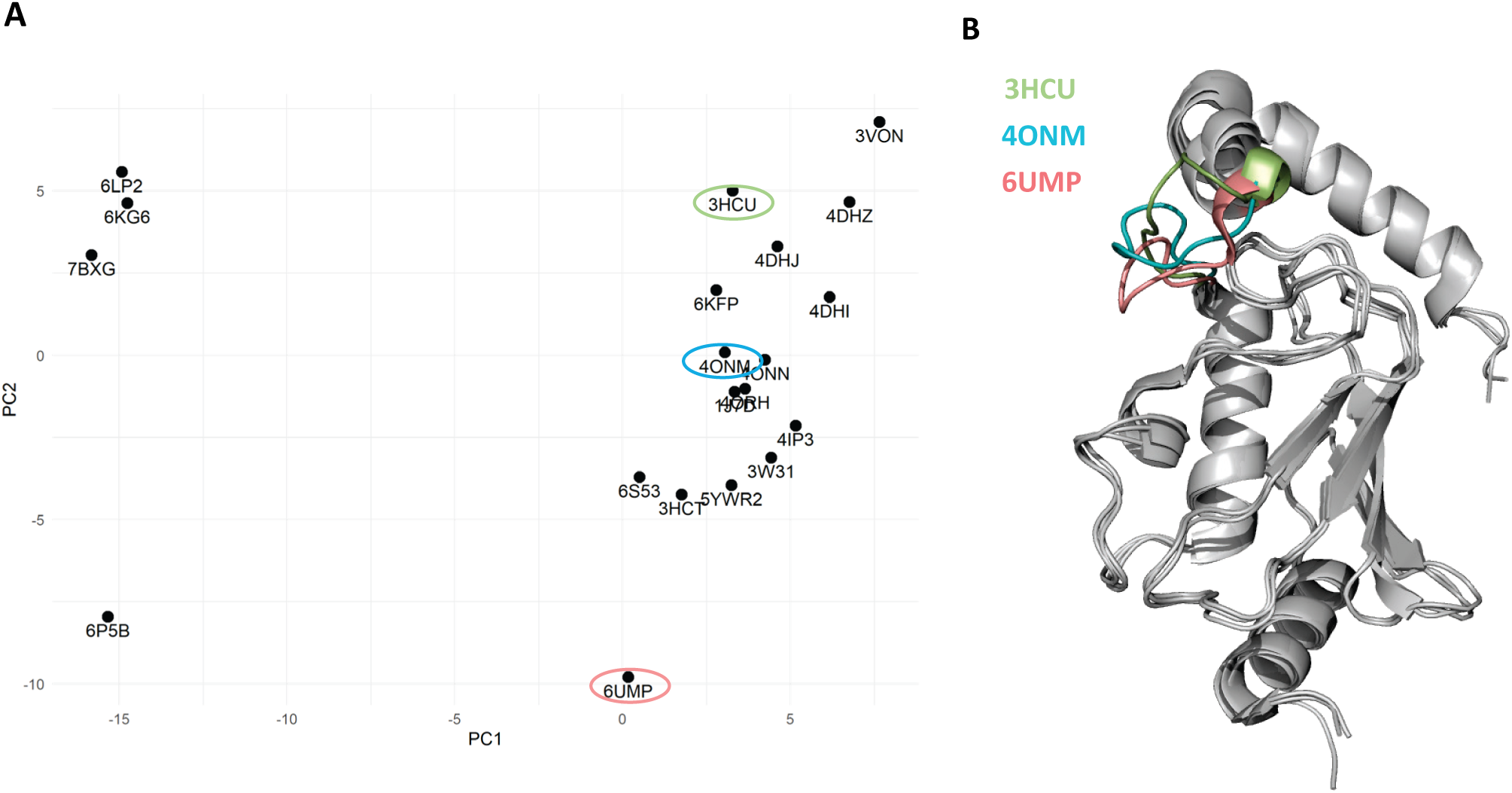
Modulation of the ubiquitin-binding site shape. **(A)** Results of the PCA analysis of UBE2N X-ray structures without mutations or gaps. **(B)** Superposition of three selected representative structures (3HCU, 4ONM, and 6UMP) illustrating differences in the ubiquitin-binding pocket of UBE2N.

To identify novel non-covalent UBE2N inhibitors, we faced the challenge of limited information regarding non-covalent binding sites on UBE2N. To address this, we performed blind docking of two known non-covalent inhibitors, ML307 and Variabine B, reported in the literature^20,21^, into the three representative UBE2N structures using AutoDockVina. During each docking run, 20 poses of ML307 and Variabine B were generated across the entire protein surface. Most poses for both inhibitors were located in two key regions considered as critical for inhibiting the ubiquitination process: the ubiquitin-binding site and the UBE2V2/UBE2V1 interaction site. We identified three distinct binding modes: binding exclusively to the ubiquitin site, exclusively to the UBE2V2/UBE2V1 site, or simultaneously to both (Figures S2, S4, and Table 1).

**Table 1.**
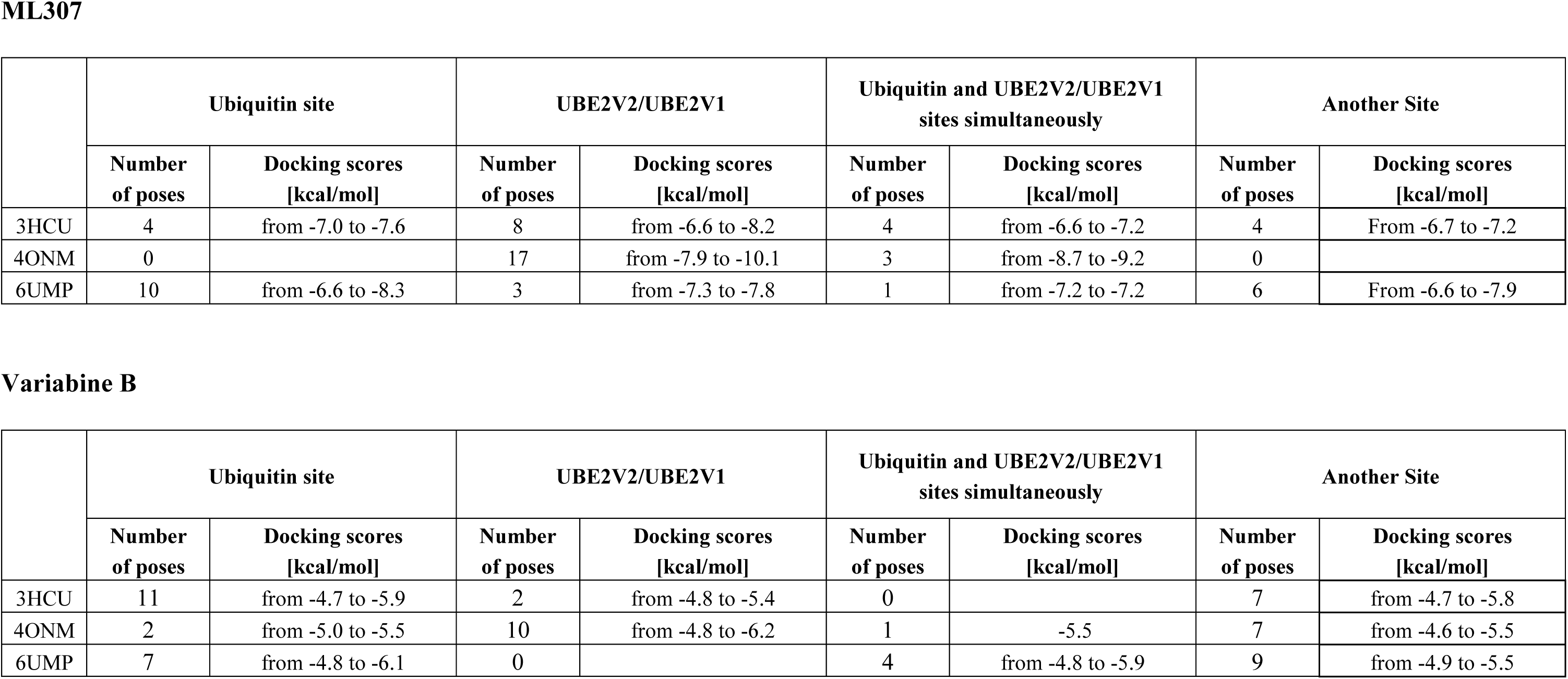
Summary of the blind docking results of ML307 and Variabine B into the three UBE2N structures.

|  | Ubiquitin site |  | UBE2V2/UBE2V1 |  | Ubiquitin and UBE2V2/UBE2V1 sites simultaneously |  | Another Site |  |
| --- | --- | --- | --- | --- | --- | --- | --- | --- |
|  | Number of poses | Docking scores [kcal/mol] | Number of poses | Docking scores [kcal/mol] | Number of poses | Docking scores [kcal/mol] | Number of poses | Docking scores [kcal/mol] |
| 3HCU | 4 | from -7.0 to -7.6 | 8 | from -6.6 to -8.2 | 4 | from -6.6 to -7.2 | 4 | From -6.7 to -7.2 |
| 4ONM | 0 |  | 17 | from -7.9 to -10.1 | 3 | from -8.7 to -9.2 | 0 |  |
| 6UMP | 10 | from -6.6 to -8.3 | 3 | from -7.3 to -7.8 | 1 | from -7.2 to -7.2 | 6 | From -6.6 to -7.9 |

**Variabine B**
|  | Ubiquitin site |  | UBE2V2/UBE2V1 |  | Ubiquitin and UBE2V2/UBE2V1 sites simultaneously |  | Another Site |  |
| --- | --- | --- | --- | --- | --- | --- | --- | --- |
|  | Number of poses | Docking scores [kcal/mol] | Number of poses | Docking scores [kcal/mol] | Number of poses | Docking scores [kcal/mol] | Number of poses | Docking scores [kcal/mol] |
| 3HCU | 11 | from -4.7 to -5.9 | 2 | from -4.8 to -5.4 | 0 |  | 7 | from -4.7 to -5.8 |
| 4ONM | 2 | from -5.0 to -5.5 | 10 | from -4.8 to -6.2 | 1 | -5.5 | 7 | from -4.6 to -5.5 |
| 6UMP | 7 | from -4.8 to -6.1 | 0 |  | 4 | from -4.8 to -5.9 | 9 | from -4.9 to -5.5 |

For ML307 docking on the 3HCU structure, 4 out of 20 poses were outside the two target sites, 4 occupied the ubiquitin site, 8 the UBE2V2/UBE2V1 site, and 4 bound simultaneously to both sites (Figure S2A, Table 1). When docked to 4ONM, all 20 poses were located in the two target regions, with 17 bound exclusively to the UBE2V2/UBE2V1 site and 3 bound simultaneously to both; no pose was found exclusively in the ubiquitin site (Figure S2B, Table 1). For 6UMP, 10 poses were located in the ubiquitin site, 3 in the UBE2V2/UBE2V1 site, and 1 bound simultaneously to both, while 6 poses were outside the target regions (Figure S2C, Table 1).

In the blind docking analysis of Variabine B, the predicted binding scores were generally less favourable compared than those of ML307 (Table 1). Interestingly, the predicted binding mode did not exclusively target the UBE2V2/UBE2V1 site, which is unexpected since previous studies suggested that Variabine B disrupts the interaction between UBE2N and its co-factors. Nonetheless, most poses were located within the two key binding sites (Figure S4). For 3HCU, 11 poses were identified in the ubiquitin site, 2 in the UBE2V2/UBE2V1 site, and 7 outside these regions. In contrast, for 4ONM, the majority of poses (10) were bound to the UBE2V2/UBE2V1 site, with 1 pose simultaneously occupying both sites, 2 in the ubiquitin site, and 7 outside the target regions (Table 1). Similarly, docking results for 6UMP showed 7 poses in the ubiquitin site, none in the UBE2V2/UBE2V1 site, 4 poses spanning both sites, and 9 poses located outside the designated regions (Figure S4C).

The blind docking results suggested that both ML307 and Variabine B have the potential to bind to ubiquitin and/or the UBE2V2/UBE2V1 sites. However, no clear preference for either compound was identified. To gain further insight, the highest-scoring poses at different sites were subjected to Molecular Dynamics (MD) refinement.

For ML307, none of the five tested complexes - two bound at the ubiquitin site, one at the cofactor site, one spanning both sites, and one bound to the OTUB1-binding site - exhibited stable binding in any of the replica simulations (SI Figure S3). Similarly, the highest-scoring Variabine B pose at the ubiquitin site proved to be unstable in all three replicas (SI Figure S5). The two highest-scoring poses at the cofactor-binding site were also subjected to MD simulations. While the highest-scoring pose was unstable, the second highest-scoring pose remained stably bound throughout 100 ns of simulation at the cofactor site in three replicas (Figure S5). In one additional replica, Variabine B left the cofactor binding site, moved around the protein, and finally rebound at another location, partially occupying the cofactor site (see Figure S5). At the starting pose producing the stable simulations, Variabine B formed an H-bond with the Asn31 side chain and established a π-stacking interaction through its aromatic core with Phe57 and Tyr34. In the three replicas, we observed that the aromatic core bound to the surface of the co-factor site and occupied the same region, but the flipping of this planar compound within the site redistributed the charged groups toward different binding spots in each replica. Energetic analysis revealed that, in this binding site, Variabine B primarily interacts through van der Waals forces, with its strongest stabilizing interaction being π-stacking with the aromatic ring of Phe57 (Figure 3). The most persistent H-bond observed across all replicas was with Glu55, occurring in approximately 23.1% in the first replica, 2.8% in the second, and 14.4% in the last. Overall, the analysis highlighted that the key residues contributing to Variabine B binding are Glu55, Phe57, Lys68 and Arg70 (Figure 3). The binding predictions at cofactor site seems more plausible as they were consistent with experimental data showing Variabine B’s ability to disrupt the UBE2N/cofactor complex.^20^

**Figure 3:**
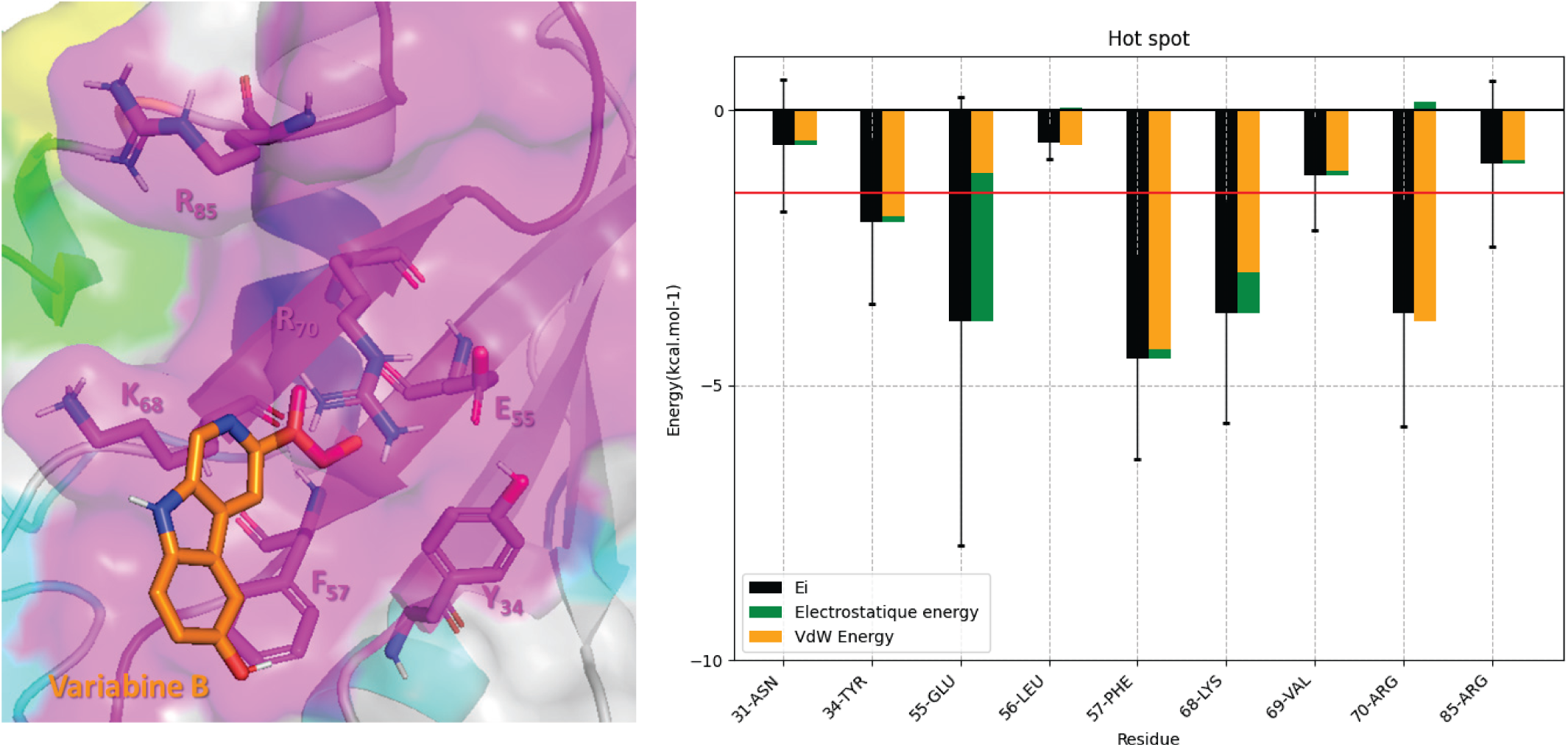
Variabine B Binding Site. Detailed view of the binding site from the final MD pose, with the side chains of interacting residues represented as sticks (Right). Only polar hydrogen atoms are shown. Interaction energies of key residues contributing to Variabine B binding (|E_INTE_| > 0.5 kcal/mol) (Left).

Despite these findings, our study provided only limited insight into the binding modes of non-covalent ligands. Notably, ML307-the most potent inhibitor reported in the literature-failed to maintain binding in any of its predicted poses.^21^ In contrast, Variabine B, described as a less potent compound, exhibited a single stable pose at the cofactor-binding site.^20^

Despite the limited information regarding the binding mode of non-covalent inhibitors on UBE2N, we initiated our screening campaign using docking techniques targeting two key regions: the ubiquitin- and the UBE2V2/UBE2V1-binding sites. The 19,000 compounds synthesized in the CERMN laboratory during previous projects and stored in our chemolibrary were converted from 2D to 3D structures, and their ionization states at pH 7.4 were predicted using OpenBabel.^31^ Only the predominant species for each compound at pH 7.4 were used in the docking screening, which was performed using AutoDockVina,^25^ separately on the two sites using the three representative structures, 3HCU, 4ONM, and 6UMP. At total 6 docking screening campaigns were carried out.

From the docking screenings, we selected a subset of promising compounds (Table 2) based on their interaction modes and their molecular size. Large compounds (> 500 Da), which are generally difficult to optimize, were excluded early in the process. The selected compounds were subsequently subjected to MD simulations. Each compound-UBE2N complex underwent a 50 ns MD simulation, during which we assessed complex stability by analysing interaction energy, ligand RMSD, and protein RMSD (calculated on Cα atoms) throughout the simulation (see example Figure S6).

**Table 2.** Summary of the Docking Screening of the CERMN chemolibrary.

|  | Ubiquitin Site |  |  |  |  |  | UBE2V2/UBE2V1 Site |  |  |  |  |  |
| --- | --- | --- | --- | --- | --- | --- | --- | --- | --- | --- | --- | --- |
|  | Docking scoring criterion [kcal/mol] | Number of selected compounds | Number of compounds submitted to MD | Number of compounds validated by MD | Number of tested compounds |  | Docking scoring criterion [kcal/mol] | Number of selected compounds | Number of compounds submitted to MD | Number of compounds validated by MD | Number of tested compounds |  |
| **3HCU** | < - 8.0 | 96 | 17 | 4 | 0 |  | < -7.5 | 54 | 4 | 0 | 0 |  |
| **4ONM** | < - 8.0 | 193 | 16 | 4 | 2 | CERMN-1<br>CERMN-6 | < -12.0 | 161 | 27 | 8 | 7 | CERMN-4<br>CERMN-5<br>CERMN-7<br>CERMN-9<br>CERMN-10<br>CERMN-12<br>CERMN-13 |
| **6UMP** | < - 8.0 | 167 | 26 | 5 | 3 | CERMN-2<br>CERMN-3<br>CERMN-11 | < -12.0 | 92 | 22 | 3 | 1 | CERMN-8 |

Only complexes exhibiting favourable stability profiles were shortlisted for further biological evaluation. In total, 24 potential inhibitors were identified *in silico*: 13 targeting the ubiquitin-binding site and 11 targeting the UBE2V2/UBE2V1 site. Due to the availability of the compounds, only 13 were selected for *in vitro* evaluation - 5 targeting the ubiquitin-binding site and 8 targeting the co-factor site.

### 3D pharmacophore screening of CERMN chemolibrary

To expand and refine our selection, we conducted an additional screening campaign using a 3D pharmacophore approach. Typically, a 3D pharmacophore is built from the structure of a reference compound bound to its target. However, since the binding modes of ML307 and Variabine B with UBE2N could not be clearly identified, we adopted alternative strategies. On one hand, we developed structure-based 3D pharmacophores by leveraging the binding site topology. On the other hand, despite the limited number of published non-covalent UBE2N,^22^ we also employed a ligand-based approach. Variabine B was chosen as template due to its lower flexibility compared to ML307, making it more suitable for pharmacophore modelling. Using LigandScout software, we generated a pharmacophore with nine pharmacophoric features based on Variabine B (Figure 4C).

**Figure 4:**
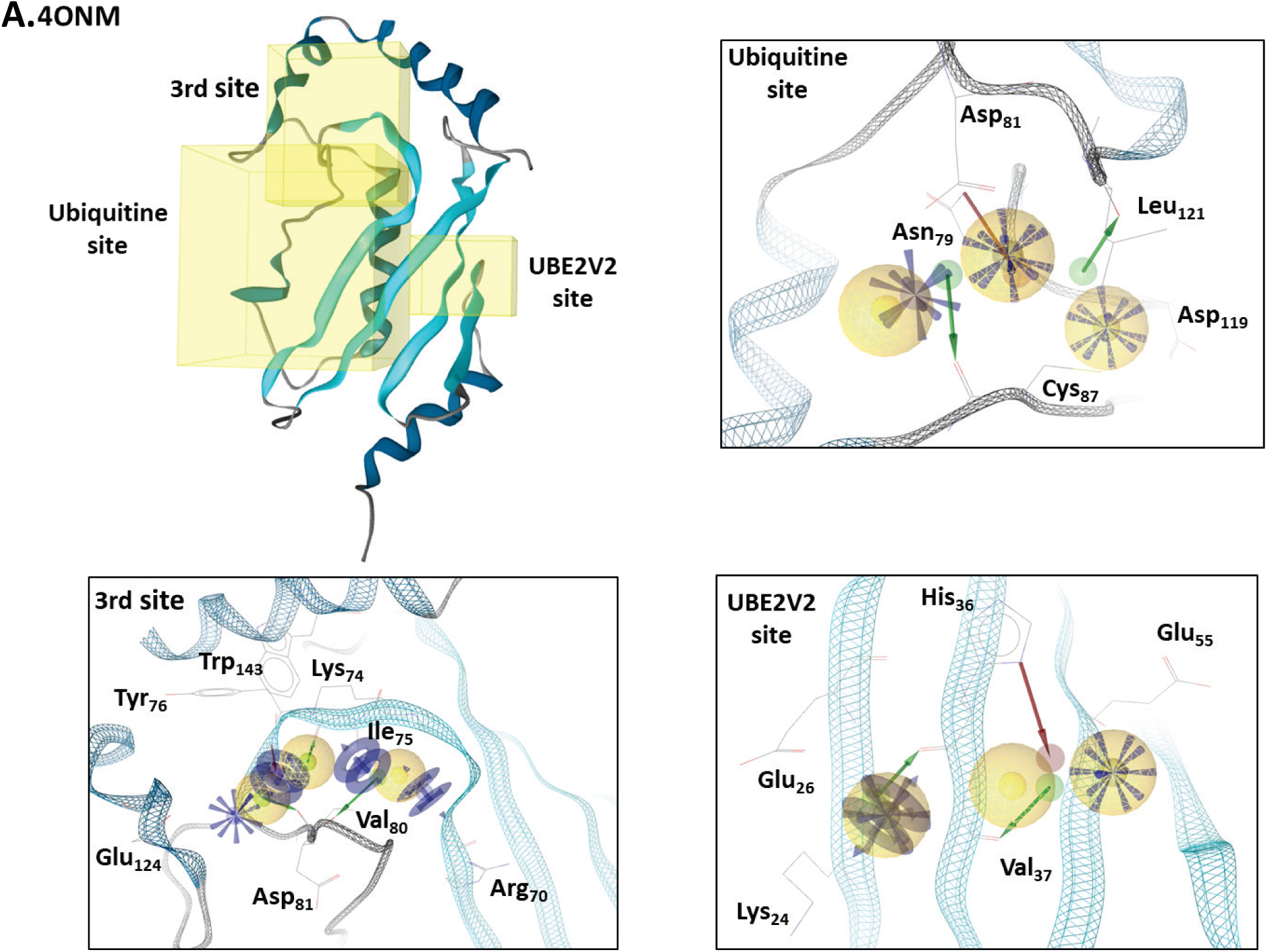
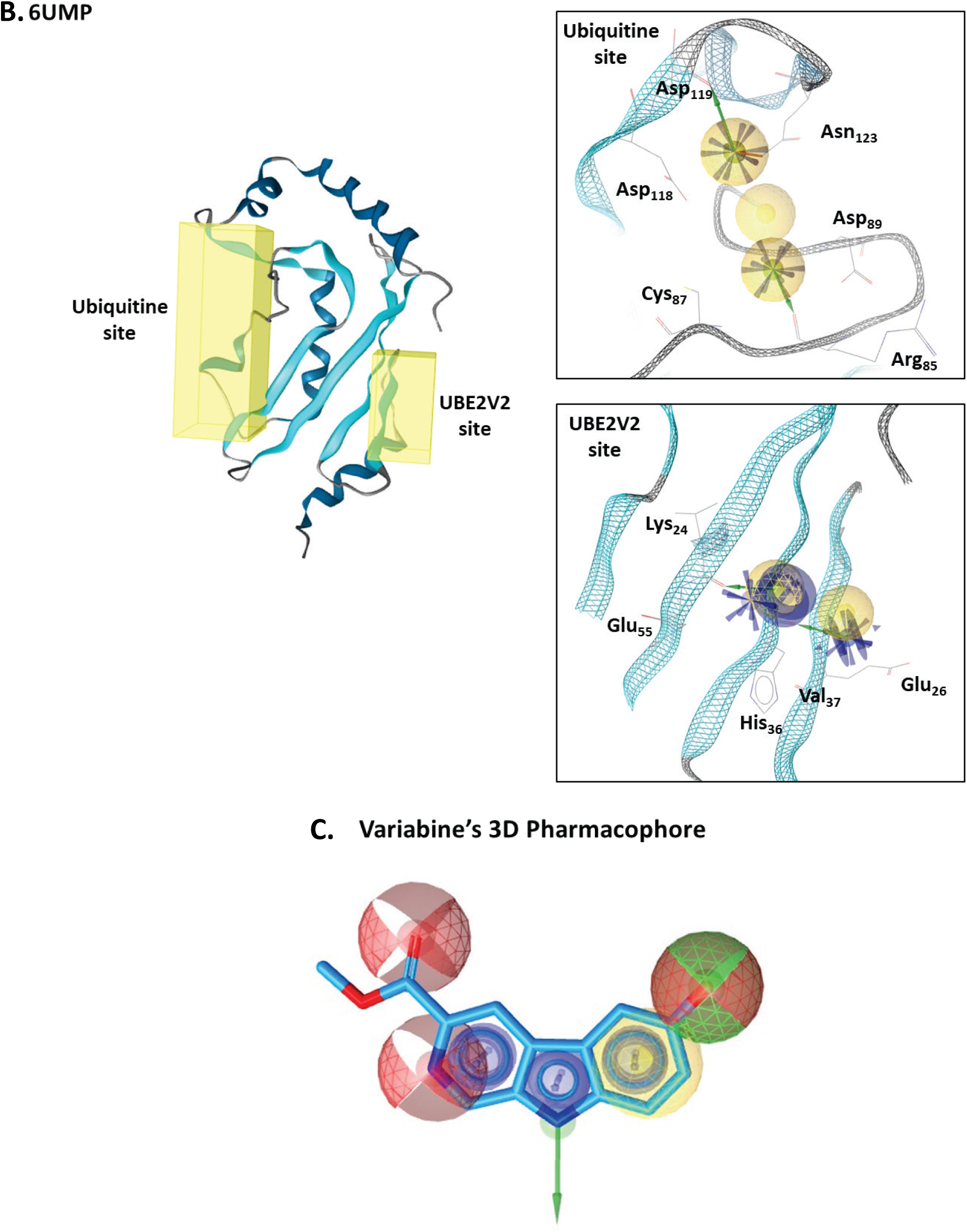
Generated 3D Pharmacophores. (A-B) The 3D pharmacophores generated from the pockets of the UBE2N X-ray structures 4ONM and 6UMP, respectively. (C) 3D pharmacophore built based on the Variabine B ligand.

To create the 3D pharmacophores from the binding site topology, we used the same three structures as for docking campaign. First, using the “Calculate Pockets” utility in LigandScout, a grid was constructed over the entire protein structure. Each grid point was evaluated based on its degree of buriedness and the number of neighbouring grid points. Isocontour surfaces were then generated, and potential binding sites were identified from them. In the case of 4ONM, LigandScout identified three binding pockets: one corresponding to the ubiquitin-binding site, a second to the UBE2V2/UBE2V1 interface, and a third partially overlapping both sites. For 6UMP, two pockets were detected, delineating the ubiquitin and UBE2V2/UBE2V1 sites. In contrast, for 3HCU, no sufficiently large pockets were identified to allow pharmacophore construction.

Next, LigandScout’s “Apo Site Grid” functionality was applied to estimate potential pharmacophore interactions at each detected site. We then used the “Apo Site Pharmacophore” tool with default settings to generate 3D pharmacophores (Figure 4A-B) which were subsequently used to screen the CERMN chemical library. The pre-processed library was loaded into LigandScout for this purpose. Since the relative importance of each pharmacophoric feature was unknown, all features were retained; however, during screening, we allowed a relatively high number of features to be omitted (for details, see Table 3).

**Table 3.**
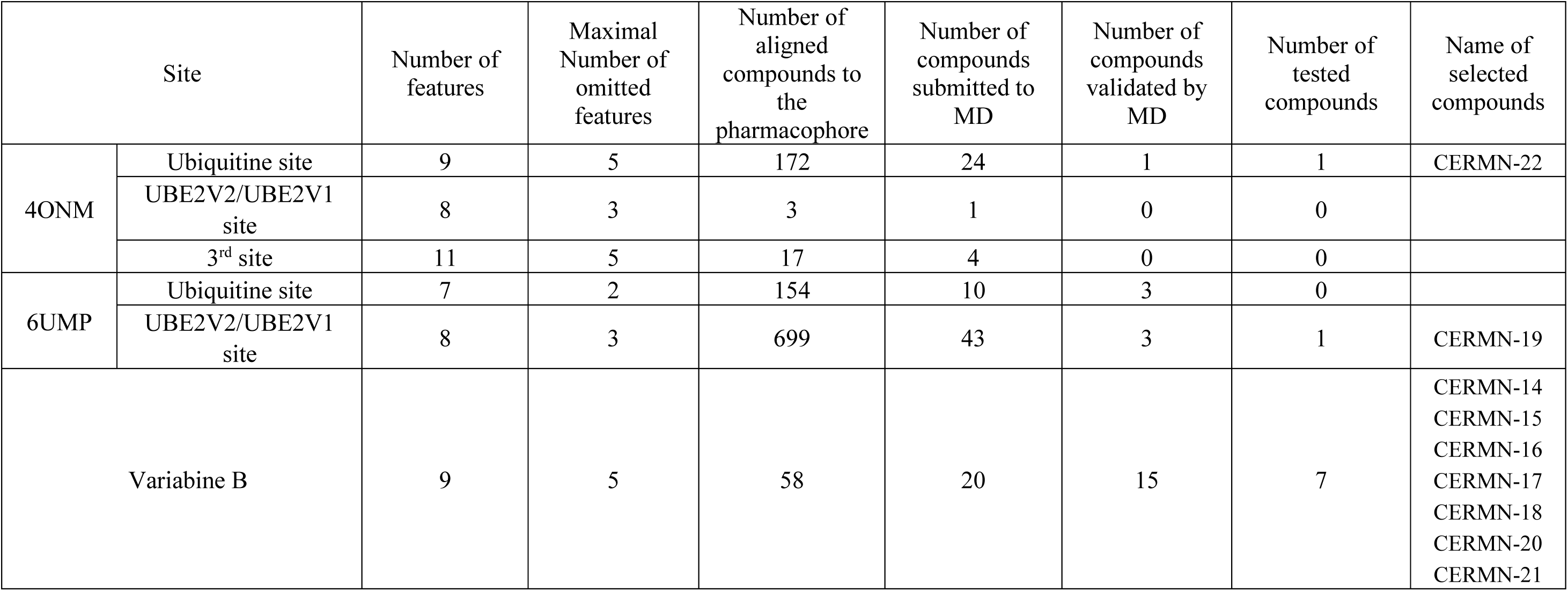
Summary of the 3D pharmacophore screening campaigns of the CERMN chemolibrary.

Aligned compounds on pharmacophore were then analysed by reviewing their docking scores from the previous docking screening campaigns. Compounds with poor docking scores, large compounds that were difficult to modulate, or those containing steroidal structures were excluded. The remaining compounds from the 3D pharmacophore screening were subjected to MD simulations to further assess their relevance and potential effectiveness, following the same approach as for the docking-based selection.

In total, 102 compound-UBE2N complexes were subjected to MD simulations, but only 22 compounds remained stably bound at the binding sites throughout the simulation. Of these, 9 compounds were selected for *in vitro* evaluation based on their availability. This selection included two compounds identified via structure-based pharmacophores: CERMN-2, predicted to target the ubiquitin-binding site, and CERMN-19, predicted to bind the cofactor site. The remaining 7 compounds, selected by the 3D pharmacophore derived from Variabine B, Variabine-like compounds, were all predicted to bind to the cofactor site (Table 3).

In total, 22 compounds were selected for *in vitro* evaluation: 13 compounds from the docking campaign and 9 compounds from the 3D pharmacophores campaign (Table 2 and 3). Of these, 6 were predicted to bind to the ubiquitin site, and 16 to the co-factor site; of which 7 were identified as Variabine-like compounds.

### Biological evaluation of compounds predicted to be UBE2N inhibitors

UBE2N inhibition has been shown to reduce cancer cell viability, as demonstrated with the validated covalent inhibitor NSC697923 and specific siRNAs.^14^ UBE2N inhibition also impairs HR DNA repair capacities^9^, hence sensitizing cancer cells to DNA damaging drugs.^14^ We thus first conducted a viability assay using the chemoresistant ovarian cancer cell line SKOV-3 to assess the effects of the compounds that we selected through our screening. Among 22 *in silico* selected compounds only 20 were tested due to solubility issues of CERMN-8 and CERMN-11. Out of these, 5 compounds were predicted to target UBE2N ubiquitin binding site, and 15 to target UBE2V1/UBE2V2 interaction site. Using a viability threshold of 60 % at a compound concentration of 40 µM, 5 out of the 20 compounds evaluated exhibited cytotoxicity. Out of the compounds targeting the ubiquitin binding site, only CERMN-2 reduced viability to 57.0 ± 7.7 % (Figure 5A). Additionally, four compounds predicted to bind the co-factor site reduced cell viability below this threshold: CERMN-16 (22.4 ± 1.7 %), CERMN-17 (52.3 ± 16.2 %), CERMN-18 (50.3 ± 4.8 %), and CERMN-20 (58.2 ± 22.8 %). Notably, compound CERMN-16 showed the strongest cytotoxic effect among all tested compounds, reducing viability to 22.4 ± 1.7% (Figure 5B). The remaining compounds had limited or no effect on SKOV-3 cells viability.

**Figure 5:**
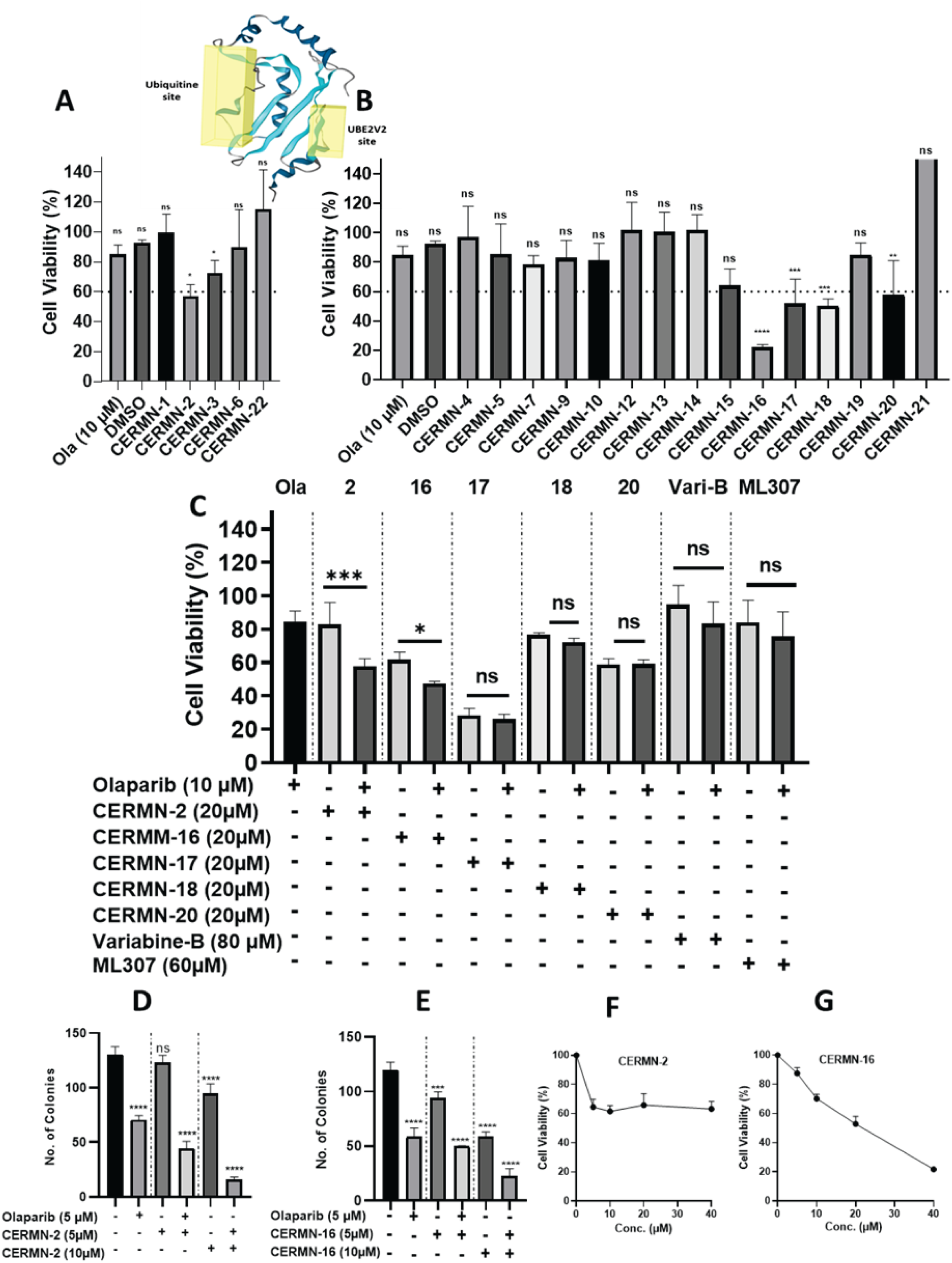
Effects of selected compounds on cancer cell and normal cell survival. (A-B) Cell viability of SKOV-3 cells treated with each of the selected 20 compounds (40 µM) using MTS assay (72 hours). (C) Sensitizing effects of five selected compounds (20 µM) on SKOV-3 to Olaparib (5 µM and 10 µM) (72 hours). (D-E) Effect of CERMN-2 and CERMN-16 on colony formation in SKOV-3 cells (5 and 10 µM), with or without Olaparib (5 µM) for 9 days. (F-G) Toxicity of CERMN-2 and CERMN-16 compounds in T1074 non-cancerous epithelial ovarian cells. Toxicity of CERMN-2 and CERMN-16 compounds in T1074 non-cancerous epithelial ovarian cells. All experiments were performed in at least three independent replicates. *p < 0.05; **p < 0.01; ***p < 0.001; ****p < 0.0001.

We selected these five compounds for further testing of their ability to sensitize SKOV3 cells to the effects of the PARP inhibitor Olaparib. At a compound concentration of 20 µM, only CERMN-2 and CERMN-16 (see SI Figure S6) significantly enhanced the cytotoxicity of Olaparib (Figure 5C). In combination with Olaparib, CERMN-2 reduced cell viability from 82.9 ±7.5 % to 57.9 ± 2.6 % and CERMN-16 from 61.7 ± 2.6 % to 47.6± 0.7 %; while Olaparib alone did show only marginal effect on cell viability at the tested concentrations (84.9 ± 3.5 %). For comparison, we also assessed three previously reported UBE2N inhibitors: NSC697923^12^, ML307^21^, and Variabine-B^20^ (synthesized in-house; see SI for detailed scheme and characterization). NSC697923 effectively sensitized SKOV3-cells and reducing viability from 76.6 ± 4.3 % to 52.3 ± 3.1 % at a concentration as low as 2 µM. In contrast, ML307 (84.3 ± 5.4 % viability) at 60 µM failed to sensitize the cells when co-treated with 10 µM Olaparib (which resulted in 75.6 ± 6.1% viability when used alone) (Figure S7A-B).

In order to validate the sensitizing effects of compounds CERMN-2 and CERMN-16, we performed a colony forming assay, again in combination with Olaparib. While Olaparib alone did show a noticeable growth inhibitory effect, the addition of CERMN-2 (10 µM) further reduced colony formation, substantially more than the compound alone (Figures 5D and S7C). Compound CERMN-16 at the same concentration produced a similar reduction in colony number when combined with Olaparib. Nevertheless, the effect appeared more additive than synergistic (Figures 5E and S7D).

Since evaluating toxicity on normal cells is essential for future pharmacological development, we tested the effects of the compounds on non-cancerous T1074 ovarian epithelial cells. NSC697923 exhibited toxicity at concentrations effective for sensitization to Olaparib (Figure S7E), while it did kill all cells at higher concentrations. In contrast, CERMN-2 was moderately toxic at sensitizing concentrations, but interestingly increasing concentrations did not exacerbate toxicity (Figure 5F). Regarding CERMN-16, it also showed moderate toxicity at effective concentrations, with increased toxicity observed at higher concentrations (Figure 5G).

### Effect of Compounds CERMN-2, CERMN-16 on UBE2N Thermal Stability

The thermal stability of the UBE2N protein was assessed using a thermal shift assay (nano-DSF), with the results presented as the derivative of fluorescent emission with respect to temperature. The melting temperature (*Tm*) determined for UBE2N alone was 66.4 ± 0.2 °C.

All compounds were evaluated in triplicate at a concentration of 100 µM (Table 4 and Figure 6). Among them, Variabine B induced the greatest stabilization, shifting the UBE2N *Tm* to 68.3 ± 0.2 °C. CERMN-2 showed a comparable thermal stabilization effect, with a *Tm* of 68.2 ± 0.3 °C. In contrast, CERMN-16 increased the protein’s *Tm* to 67.1 ± 0.2 °C, only slightly above the control value (66.4 ± 0.2 °C), indicating a weaker stabilizing effect than that observed for Variabine B or CERMN-2.

**Figure 6:**
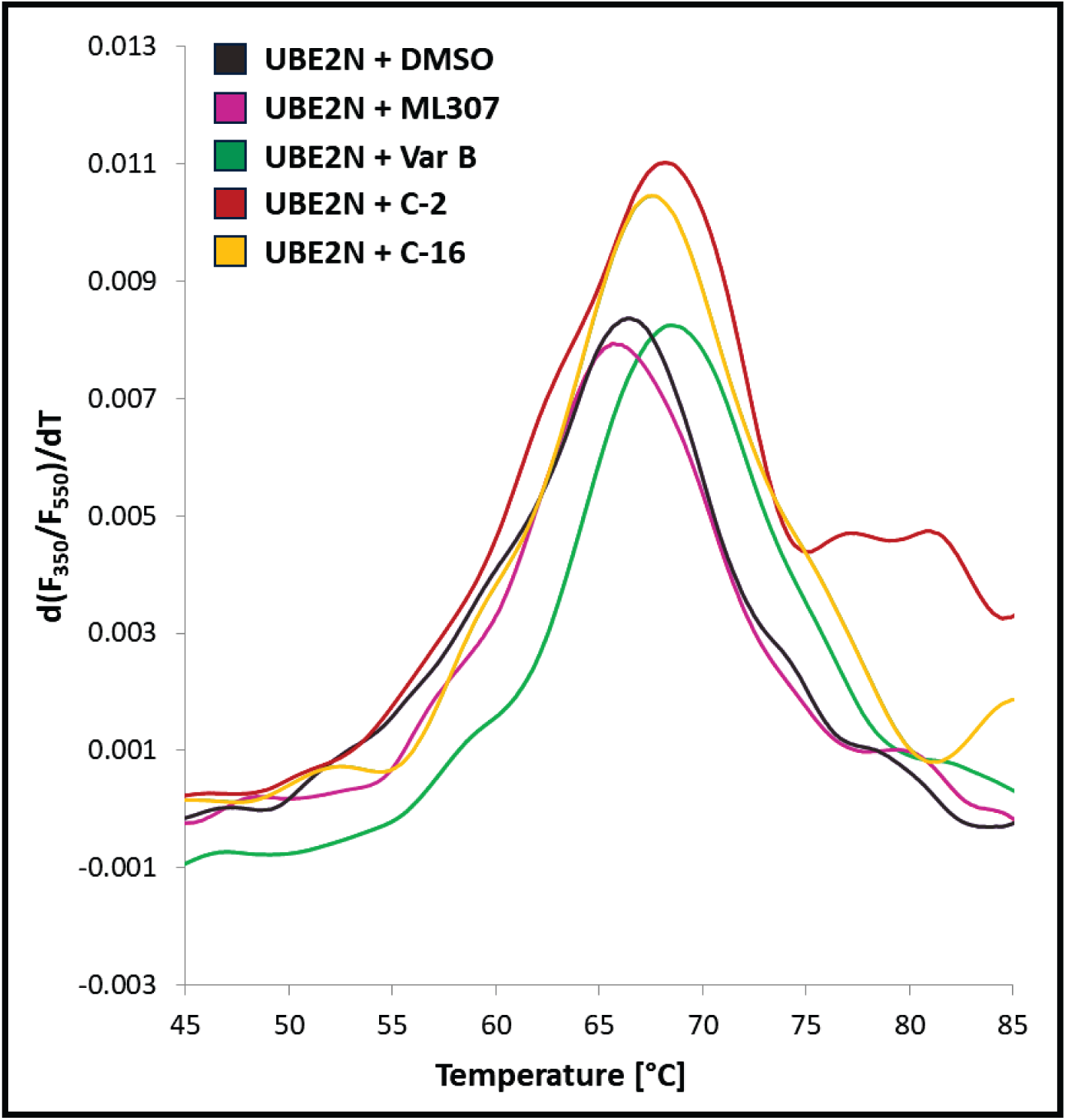
Impact of compounds on the thermal stability of UBE2N measured by nanoDSF. The results are presented as a derivative of fluorescent emission with respect to temperature, showing a representative curve from one of three independent experiments. The melting temperature (*Tm*) of UBE2N was determined in the absence of inhibitor (black, *Tm* = 66.4 ± 0.2 °C) and in the presence of 100 µM of each compound: ML307 (pink, *Tm* = 65.8 ± 0.1 °C); Variabine B (green, *Tm* = 68.3 ± 0.2 °C); CERMN-2 (red, *Tm* = 68.2 ± 0.3 °C); CERMN-16 (yellow, *Tm* = 67.1 ± 0.2 °C).

**Table 4.** Melting Temperatures (*Tm*) of UBE2N Determined by Thermal Stability Assay in the presence of potential UBE2N inhibitors.

| **Condition/**<br>**Compound** | **Melting Temperature**<br>**($T_m$) (°C)** |
| --- | --- |
| UBE2N (No inhibitor) | $66.4 \pm 0.2$ |
| CERMN-2 | $68.2 \pm 0.3$ |
| CERMN-16 | $67.1 \pm 0.2$ |
| Variabine B | $68.3 \pm 0.2$ |
| NSC | $67.7 \pm 0.3$ |
| ML307 | $65.8 \pm 0.1$ |

ML307 exhibited no stabilizing capacity toward UBE2N and showed the lowest melting temperature (65.8 ± 0.1 °C), below that of UBE2N alone. This suggests that ML307 acts as a destabilizer rather than an inefficient stabilizer of the protein structure.

### Binding Affinity of CERMN-2 to UBE2N by Microscale Thermophoresis (MST)

Microscale Thermophoresis (MST) was employed to determine the binding affinity (*K_d_*) of selected compounds to UBE2N. Variabine B was found to bind to UBE2N with a dissociation constant (*K_d_*) of 35 µM, which is consistent with previously reported values by Sakai *et al.* (16 µM).^20^ More precisely, dose-response curves derived from MST traces recorded at 9-10 seconds MST-on time (n = 3) depict the binding of Variabine B to UBE2N, yielding a *K_d_* of 35.3 ± 6.3 µM. Although the literature reports that ML307,^21^ directly is bound to UBE2N, our MST data do not support its designation as a “known potent non-covalent inhibitor” under the conditions assessed. Among the two discovered candidates, CERMN-2 exhibited a slightly lower affinity for UBE2N compared to Variabine B, with a *K_d_* of 57.7 ± 3.1 µM. In contrast, CERMN-16 showed no detectable binding to UBE2N under the tested MST conditions. Figure 7 presents the binding curves illustrating the direct interaction between UBE2N and the compounds Variabine B and CERMN-2. The binding data obtained here correlate well with the thermal stability results, in which the two confirmed binders, Variabine B and CERMN-2, produced the greatest thermostabilization of UBE2N (Tm of 68.3 ± 0.2 °C and 68.2 ± 0.3 °C, respectively, at 100 µM).

**Figure 7:**
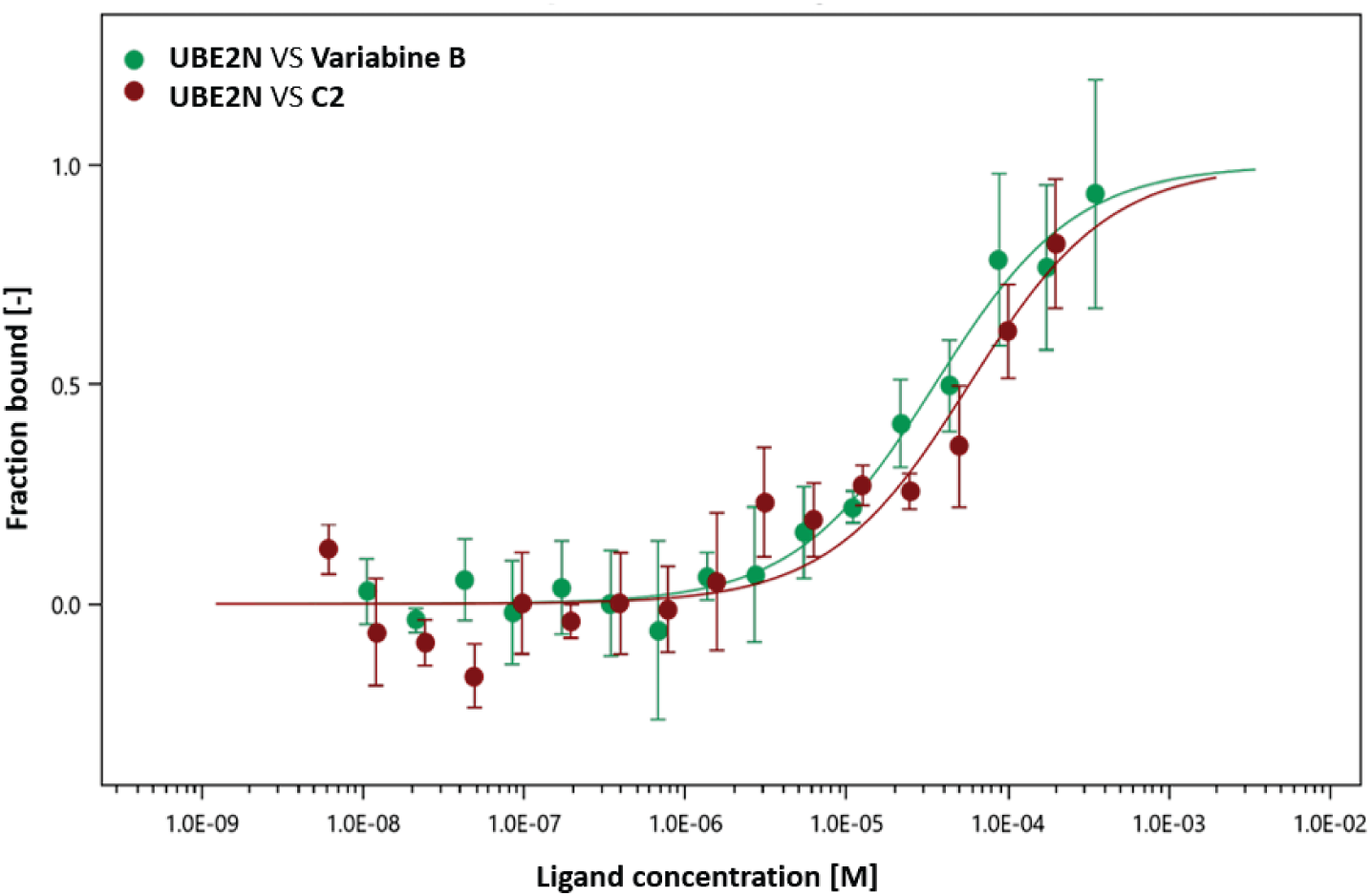
MST binding assay showing the interaction of Variabine B and CERMN-2 with UBE2N. Dose-response curves depict the binding of Variabine B (green, *K_d_* = 35.3 ± 6.3 µM) and CERMN-2 (red, *K_d_* = 57.7 ± 3.1 µM) to UBE2N, derived from MST traces recorded at 9–10 s MST-on time (n = 3).

### In depth in-silico prediction of CERMN-2 binding mode

Among the 20 compounds evaluated *in vitro*, CERMN-2 demonstrated the most promising activity and specific binding to UBE2N, warranting a more in-depth analysis of its binding modes. CERMN-2, identified through a docking campaign using the 6UMP structure, achieved a docking score of -8.3 kcal/mol and was predicted to bind at the ubiquitin-binding site.

Docking studies predicted that CERMN-2 binds at the ubiquitin site, near Lys68. In the docked pose, no hydrogen bonds were formed with UBE2N. However, one of CERMN-2’s aromatic rings was positioned close to the protonated side chain of Lys68 (Figure S6). Although the ligand’s movement within the cavity differed slightly among the replica simulations, the overall displacement pattern was the same: the ligand shifted to the opposite side of the cavity, toward the flexible loop between residues 114–124.

In the first replica, the displacement occurred around 50 ns; the ligand then rotated to better accommodate this region of the binding site, where it remained stable for the next 50 ns. In the other two replicas, the shift of the ligand occurred early, within the first few nanoseconds of the simulation. In the second replica, the ligand then rotated within the binding site before stabilizing at around 50 ns, whereas in the third replica the stable position was reached very quickly (see Figure S8). The aromatic rings in the stable poses obtained from the three simulations occupied the same cavity; the molecule simply flipped by 180°, resulting in one pose in which the NC=OH (amide) group pointed toward Cys87, while in the other two it pointed in the opposite direction. In all three simulations, the generated poses obstruct access to the catalytic Cys87 (Figures 8 and S8).

**Figure 8:**
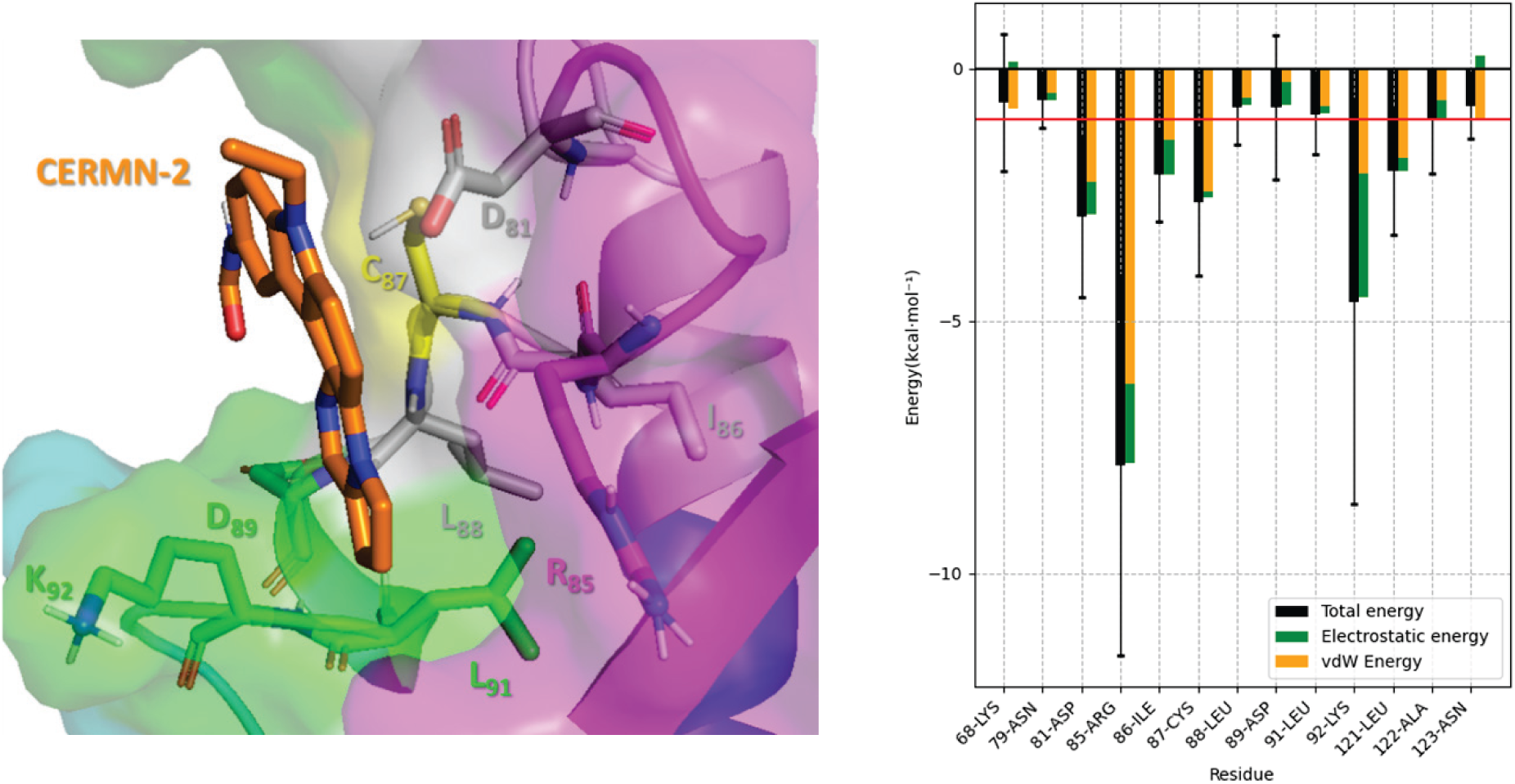
CERMN-2 predicted binding mode. Representative view of the CERMN-2 binding site from the MD replica, with the side chains of interacting residues represented as sticks. Only polar hydrogen atoms are shown (Right). Interaction energies of key residues contributing to CERMN-2 binding (|E_INTE_| > 0.5 kcal/mol) (Left).

Across all replicas, changes in the ligand position correlated with fluctuations of the flexible loop, which dynamically altered the shape of the cavity. Although both positively and negatively charged residues contribute to the ubiquitin-binding site, no persistent hydrogen bond (i.e., present in more than 25% of the MD trajectory) was observed in any replica. The most frequent hydrogen bonds were formed with the Lys92 side chain (6.0% in the first replica, 14.6% in the second, and 22.6% in the third). Energetic analysis confirmed that electrostatic interactions contribute less than van der Waals interactions, indicating that van der Waals forces are the primary driving force for CERMN-2 binding (Figure 8). The residues predicted to contribute most to CERMN-2 binding were Arg85, Lys92, Asp81, and Cys87. Indeed, our simulations predict that CERMN-2 binds to the ubiquitin-binding site due to its structural complementarity with the cavity, thereby effectively blocking access to the catalytic Cys87 residue.

In conclusion, our molecular modelling studies support CERMN-2 as a genuine inhibitor of ubiquitin binding, acting similarly to a covalent inhibitor. Indeed, our results suggest that CERMN-2 obstructs access to the catalytic Cys87.

## DISCUSSION

UBE2N has emerged as a promising therapeutic target for cancer due to its critical roles in DNA repair and NF-κB signaling pathways.^9^ While covalent inhibitors targeting the catalytic Cys87 residue have shown preclinical potential, non-covalent inhibitors offer greater promise, particularly in terms of selectivity and reduced off-target effects. Several non-covalent inhibitors, annotated as disrupting the UBE2N/UBE2V1 interaction, have been previously reported - most of which are natural compounds.

Leucettamol A, isolated from *Leucetta aff. microraphis*, was the first reported disruptor of the UBE2N/UBE2V1 interaction, with an IC_50_ of approximately 50 µg/mL (equivalent to 106 µM) in an ELISA assay.^18,21^ However, its potential is limited by instability and a propensity for isomerization. Notably, another study reported that Leucettamol A showed no detectable activity against UBE2N.^21^ More recently, Manadosterols A and B, isolated from *Lissodendryx fibrosa*, were identified as significantly more potent inhibitors of the UBE2N/UBE2V1 interaction, with IC_50_ values of 0.09 µM and 0.13 µM, respectively, as determined by ELISA.^19^ These compounds, however, were not further investigated. Variabine B, another natural inhibitor reported to disrupt the interaction between UBE2N and its cofactors UBE2V1 or UBE2V2, is a β-carboline alkaloid and was reported to inhibit the UBE2N/UBE2V1 interaction with an IC_50_ of 4 µg/mL (equivalent to 16 µM).^20^

Among synthetic non-covalent inhibitors, ML307 has been described as a potent sub-micromolar UBE2N inhibitor, although it remains unclear whether it targets the ubiquitin-binding site, the cofactor interface, or another site. It exhibited an IC_50_ of 0.781 µM in a TR-FRET-based ubiquitination assay and was considered the most potent UBE2N inhibitor tested *in vitro* at the time.^21^ Despite favourable solubility, stability in buffer and plasma, and selectivity against tested targets, ML307 suffered from poor microsomal stability, limiting its translational potential.

The literature lacks structural information describing the molecular binding modes of reported non-covalent UBE2N inhibitors. To address this gap, we conducted a study to determine the binding modes of ML307 and Variabine B. Molecular docking of ML307 revealed limitations in predicting stable binding: despite favourable docking scores, it failed to maintain a stable pose during MD simulations. In contrast, Variabine B exhibited at least one stable pose in MD simulations but showed lower cellular activity than ML307 in SKOV-3 cells (Figure S7).

To address the need for novel, well-characterized non-covalent inhibitors, we conducted a rational *in silico* screening strategy integrating molecular docking and 3D pharmacophore modelling, targeting two potential binding sites: the ubiquitin-binding site and the cofactor interface. Additionally, supplementary 3D pharmacophore was generated based on the chemical structure of Variabine B. Multiple screening campaigns using of a library of approximately 19,000 compounds - systematically validated by MD simulations to confirm ligand binding stability - ultimately led to the selection of 22 candidates for biological evaluation, of which only 20 could be tested due to solubility issues. Distinct hit rates were observed for each screening method. Molecular docking yielded 13 compounds, of which 11 were biologically evaluated - 4 targeting the ubiquitin-binding site and 7 targeting the co-factor binding site. Among these, only one compound (CERMN-2) reduced the viability of the chemoresistant ovarian cancer cell line SKOV-3 by more than 40%. In parallel, 3D pharmacophore modelling based on the topology of the binding sites identified two additional compounds compared to the docking approach - one targeting the ubiquitin site and one the co-factor site - but neither demonstrated a significant effect on cell viability. Taken together, this corresponds to a success rate of approximately 8% for the structure-based screening strategy.

In contrast, seven compounds were selected using a 3D pharmacophore model derived directly from the structure of the known ligand Variabine B, representing a ligand-based drug design approach. Of these, four compounds reduced SKOV-3 cell viability by more than 40% - CERMN-16; CERMN-17, CERMN-18 and CERMN-20 - corresponding to a success rate of approximately 57%. These results highlight the differing efficiencies of the two strategies. Although structure-based screening showed a lower hit rate compared to the ligand-based approach, it holds the potential to identify structurally novel inhibitor that differ from those previously reported.

The most potent compound identified through ligand-based 3D pharmacophore screening, CERMN-16, was predicted to mimic the binding mode of Variabine B and thus target the UBE2N cofactor interface. However, CERMN-16 only marginally affected UBE2N thermal stability, and did not bind to UBE2N in our MST assay, contrasting with Variabine B, which we showed to interact with UBE2N, as well as affecting its thermal stability. While CERMN-16 unsurprisingly does not seem to genuinely sensitize SKOV-3 cells to Olaparib according to interaction results, the absence of sensitization by Variabine B is unexpected. The predicted binding mode of these two compounds, at the cofactor interface, contrasts with that of covalent inhibitors, which target the catalytic Cys87 residue and occupy an adjacent cavity that is largely inaccessible to non-covalent ligands. We can therefore hypothesize that despite binding to free UBE2N, Variabine B may not be able to displace UBE2N from its interaction with its cofactors, or outcompete them, *in vivo* in cells.

The lack of cellular sensitization by ML307, even at high concentration (60 µM), is particularly striking given its reported submicromolar *in vitro* inhibition of UBE2N (IC_50_ = 781 nM) and its favourable pharmacokinetic properties, including good aqueous solubility, plasma stability, and transcellular permeability. However, these *in vitro* and pharmacokinetic advantages did not translate into a thermal shift nor into a sensitization effect with Olaparib in the cellular context.

In contrast, CERMN-2 identified through structure-based docking, was predicted to bind at the ubiquitin-binding site. In addition, we could show that its presence modified the thermal stability of UBE2N, and that it could bind to free UBE2N in our MST experiments. MD simulations suggested that van der Waals interactions are the main contributors to its binding affinity, and that the compound effectively blocks access to the catalytic Cys87 residue by occluding the ubiquitin interaction interface. This finding highlights a potentially unique mechanism and provides new insights into non-covalent inhibition at the ubiquitin-binding site of UBE2N, a region for which structural or modelling data on non-covalent inhibitors is currently lacking in the literature.

Taken together, our results highlight CERMN-2 as a promising direct non-covalent inhibitors of UBE2N, a key target in cancer research.^14,22^ Future studies will be needed to explore its ability to interfere with UBE2N’s role in DNA repair and to sensitize to DNA damaging drugs more relevant study models, such as patient-derived tumour organoids of ovarian cancer.^47^

## CONCLUSION

In this study, we addressed the need for novel non-covalent UBE2N inhibitors by applying a dual *in silico* approach combining structure-based and ligand-based strategies. Among the 20 tested compounds, CERMN-2 emerged as the most promising direct non-covalent inhibitors of UBE2N. CERMN-2, identified *via* structure-based screening, also demonstrated sensitizing properties and a favourable safety profile in normal ovarian epithelial cells. Molecular dynamics simulations provided insights into its distinct binding modes at the ubiquitin-binding site, further supported by its ability to perturb thermal stability of UBE2N and directly bind to the protein. These results highlight the therapeutic potential of CERMN-2 as non-covalent UBE2N inhibitor. Although further validation of direct target engagement is needed, our findings pave the way for the development of a new generation of selective, non-natural UBE2N inhibitors for combination cancer therapy.

## ASSOCIATED CONTENT

**Supporting Information**. (Table S1) Summary of box parameters in Å applied in the blind docking studies and the docking screening; (Table S2) The applied Threshold and Buriedness values in Pocket Detection functionality of the LigandScout; (Figure S1) Results of PCA analysis. (Figure S2) The results of the blind docking of ML307 on the three selected UBE2N X-ray structures selected for the docking study; (Figure S3) The results of the molecular dynamics simulations on the selected poses of ML307 at the ubiquitin site, the UBEV2/UBE2V1 site and OTUB1 binding site; (Figure S4) The results of the blind docking of Variabine B on the three selected UBE2N X-ray structures selected for the docking study; (Figure S5) The results of the molecular dynamics simulations on the selected poses of Variabine B in the ubiquitin site and in the UBEV2/UBE2V1 site; (Figure S6) Example of the applied molecular screening strategy; (Figure S7) Effects of Covalent and Non-Covalent UBE2N Inhibitors Alone or Combined with Olaparib on SKOV3 Cell Proliferation, Clonogenicity, and Toxicity; (Figure S8) The replica of the molecular dynamics simulations on the selected pose of CERMN-2 in the ubiquitin site; Synthesis of Variabine B.

This material is available free of charge via the Internet at http://pubs.acs.org.

## Author Contributions

‡ Côme Ghadi and Shafi Ullah Khan contributed equally. The manuscript was written through contributions of all authors. All authors have given approval to the final version of the manuscript.

## ACKNOWLEDGMENT

We thank the CRIANN (Centre Régional Informatique et d’Applications Numériques de Normandie) for the computing facilities.

## DATA AND SOFTWARE AVAILABILITY

LigandScout 4.4.9 is a software developed and distributed by Inte:Ligand GmbH (www.inteligand.com/ligandscout). Requests for access should be addressed directly to the company.

All other software used in this study are open-source (OpenBabel, CHARMM, NAMD, AutoDock Vina, ChemAxon, CHARMM-GUI – htpps://charmm-gui.org), and the input data required to reproduce the study are provided in the accompanying ZIP file: SI_docking complexes and pharmacophores.zip.

