## Supporting Information for "Integrative In Silico and Experimental Identification of Non-Covalent UBE2N Inhibitors Enhancing PARP Inhibitor Sensitivity"

[¥] Univ. Lille, CHU Lille, CNRS, Inserm, UMR9020 - UMR1277 - Canther - Cancer Heterogeneity, Plasticity and Resistance to Therapies, F-59000 Lille, France.

[£] Univ. Lille, CHU Lille, ULR 7365 - GRITA - Groupe de Recherche sur les formes Injectables et les Technologies Associées, F-59000 Lille, France.

* Co-corresponding authors

**Table S1.** Summary of box parameters in Å applied in the blind docking studies and the docking screening

| Docking | PDB | X Size | Y Size | Z Size | X Center | Y Center | Z Center |
|---|---|---|---|---|---|---|---|
| ML307 Blind Docking | 3HCU | 44 | 48 | 42 | 32.15 | 24.027 | 35.063 |
| | 4ONM | 44 | 52 | 48 | 32.158 | 22.738 | 36.19 |
| | 6UMP | 40 | 42 | 48 | 34.081 | 18.994 | 33.853 |
| Variabine Blind Docking | 3HCU | 42 | 48 | 42 | 32.162 | 24.809 | 34.254 |
| | 4ONM | 46 | 48 | 48 | 32.167 | 22.972 | 34.254 |
| | 6UMP | 44 | 50 | 46 | 32.378 | 23.886 | 33.503 |
| Screening on Ubiquitin site | 3HCU | 20 | 20 | 10 | 38.044 | 18.534 | 26.796 |
| | 4ONM | 20 | 10 | 20 | 38.416 | 18.14 | 27.165 |
| | 6UMP | 12 | 22 | 18 | 29.506 | 18.994 | 33.853 |
| Screening on UBE2V2 site | 3HCU | 20 | 20 | 20 | 22.812 | 19.371 | 32.184 |
| | 4ONM | 18 | 10 | 14 | 25.924 | 17.14 | 33.226 |
| | 6UMP | 20 | 16 | 14 | 39.755 | 18.855 | 24.93 |

**Table S2.** The applied Threshold and Buriedness values in Pocket Detection functionality of the LigandScout

| | 3HCU | 4ONM | | | 6UMP | |
|---|---|---|---|---|---|---|
| | Site1 | Site1 | Site2 | Site3 | Site1 | Site2 |
| Buriedness | 0 | 0.21 | 0.1 | 0 | 0.2 | 0.1 |
| Threshold | 0.3 | 0.3 | 0.3 | 0.3 | 0.3 | 0.3 |

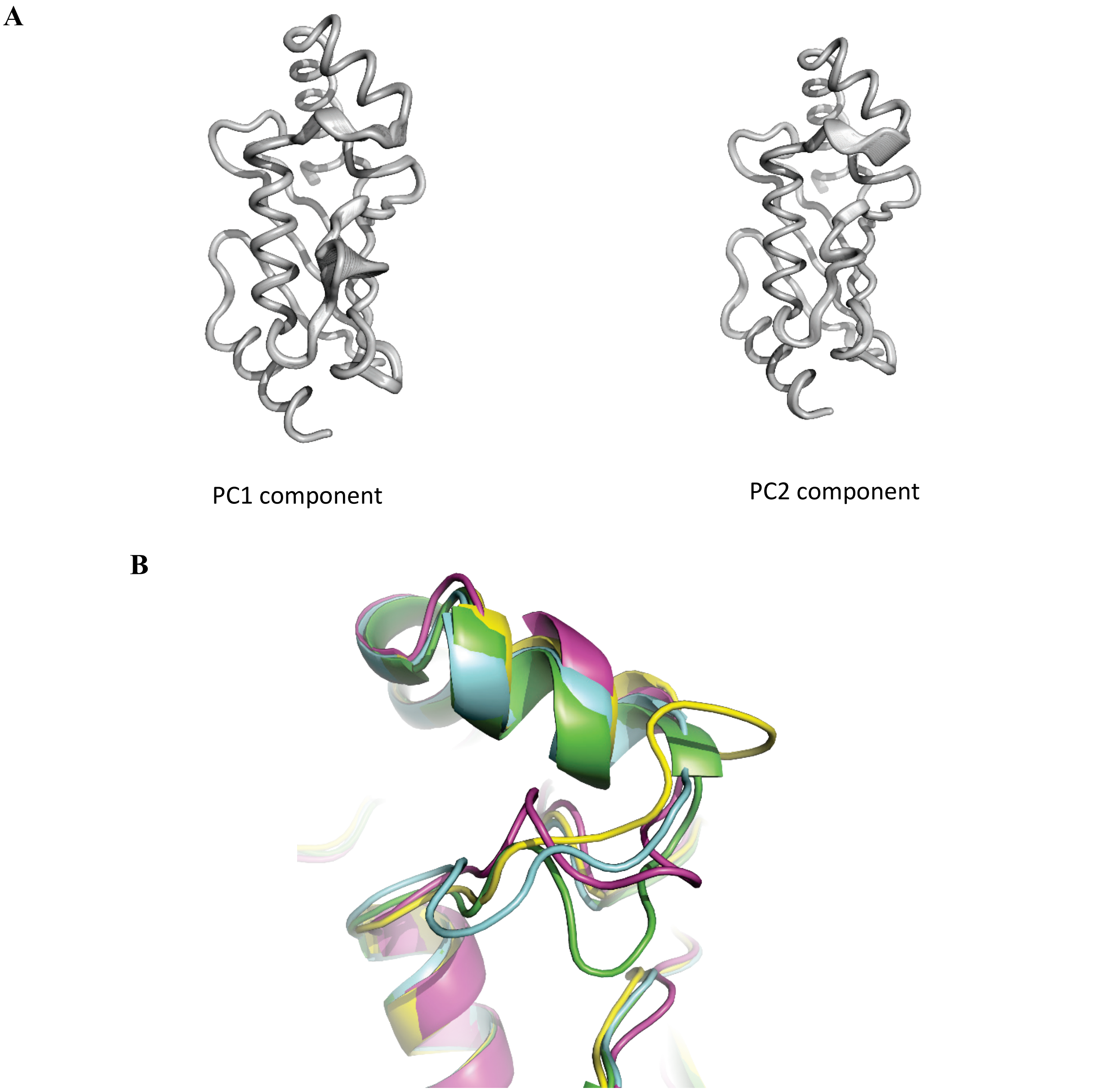


**Figure S1.** Results of PCA analysis. (A) Tube representation of the PC1 and PC2 principal components. (B) Superposition of the structures with PDB ID: 3VON (cyan), 3HCU (green), 4ONM (pink) and 6UMP (yellow).

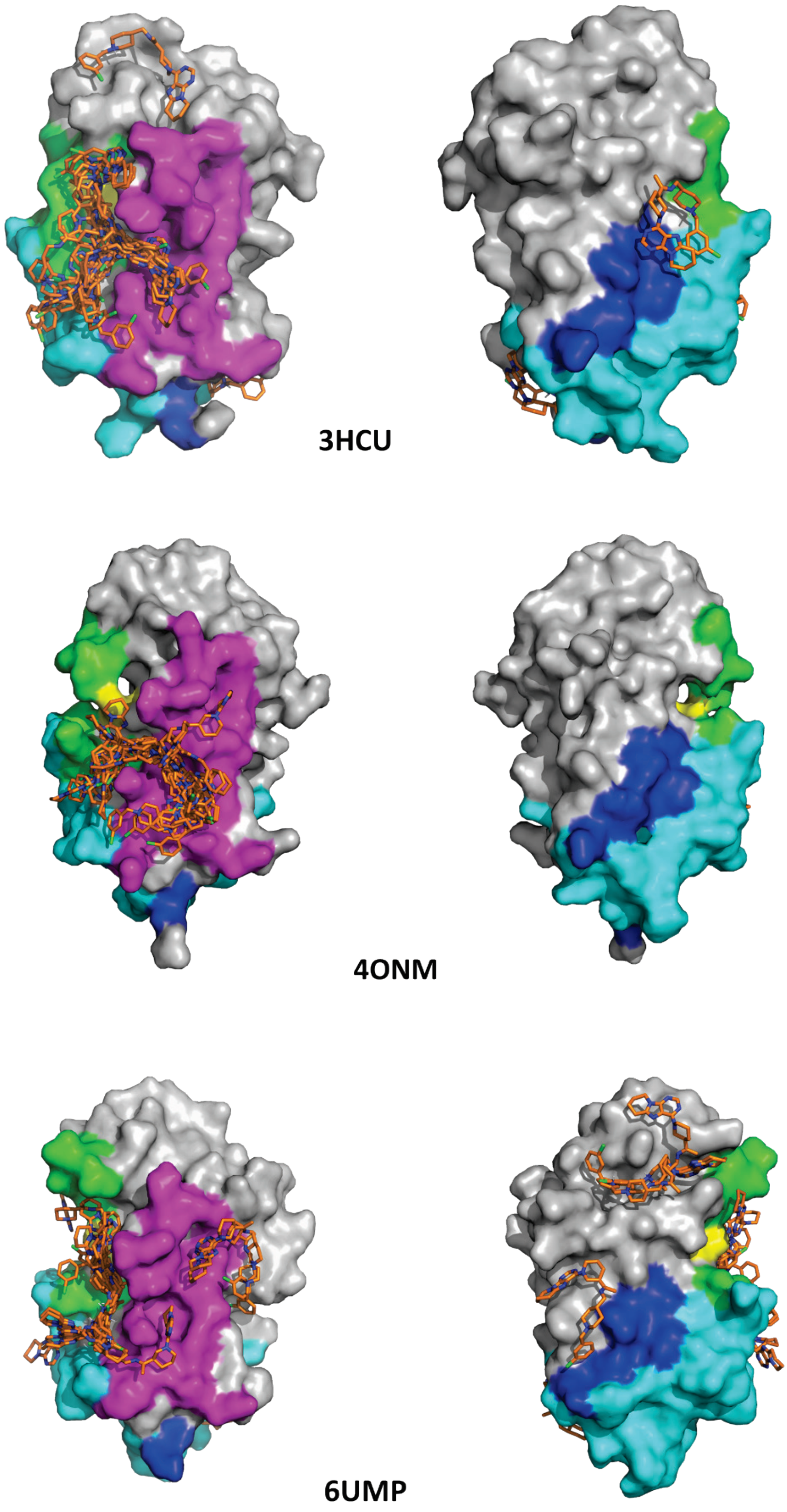


**Figure S2:** The results of the blind docking of ML307 on the three selected UBE2N X-ray structures selected for the docking study.

#### ML307: 6UMP ubiquitin site – pose 1

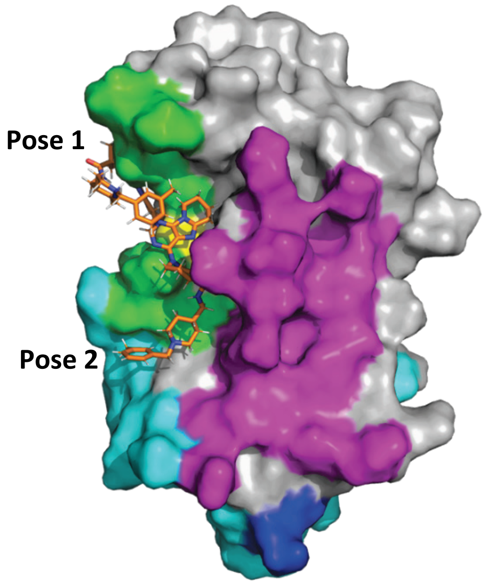


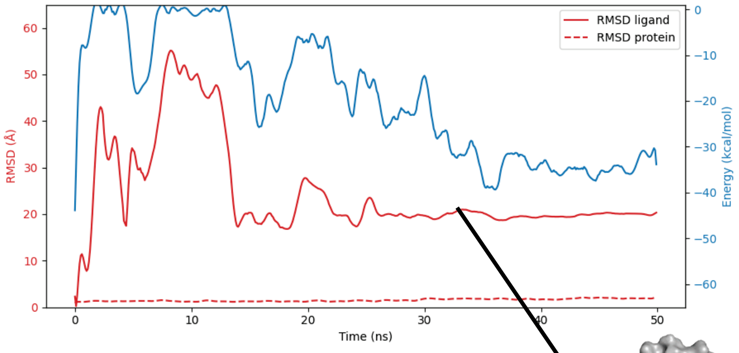


Stabilisation of the ligand in another site

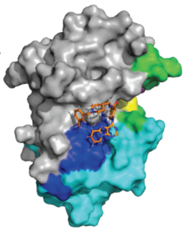

Leave of the ligand from ubiquitin site

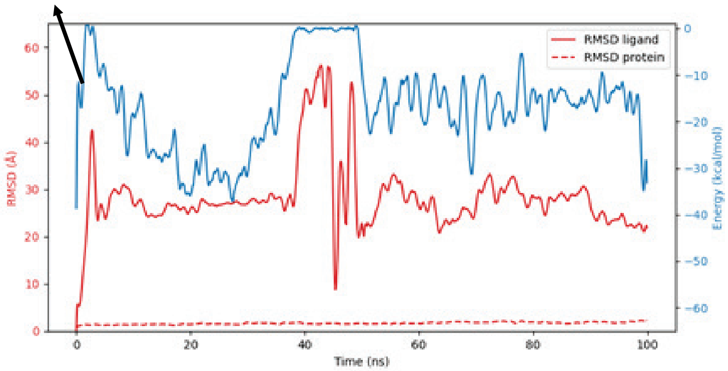


Leave of the ligand from ubiquitin site

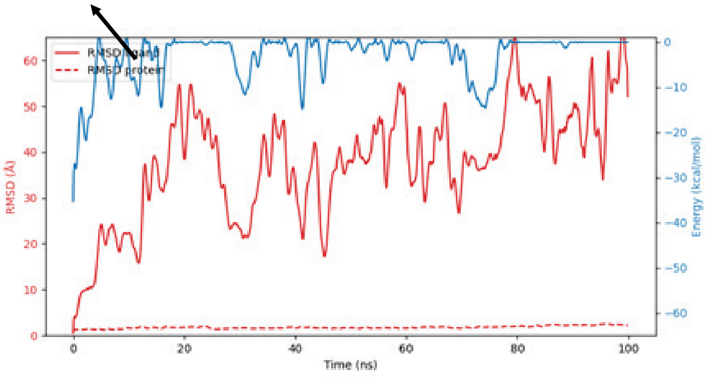

### ML307: 6UMP ubiquitin site – pose 2

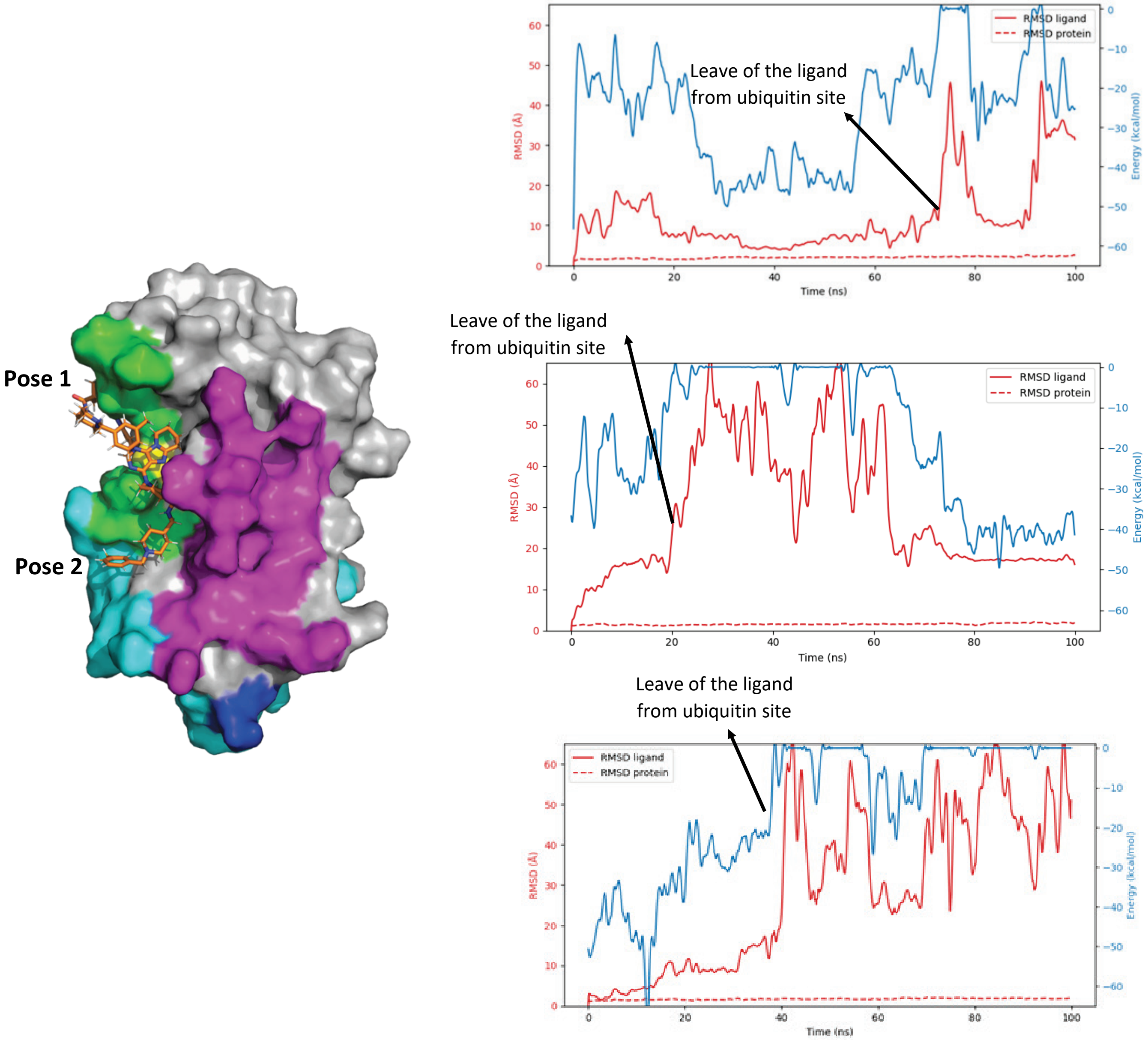

### ML307: 4ONM ubiquitin site – pose 1

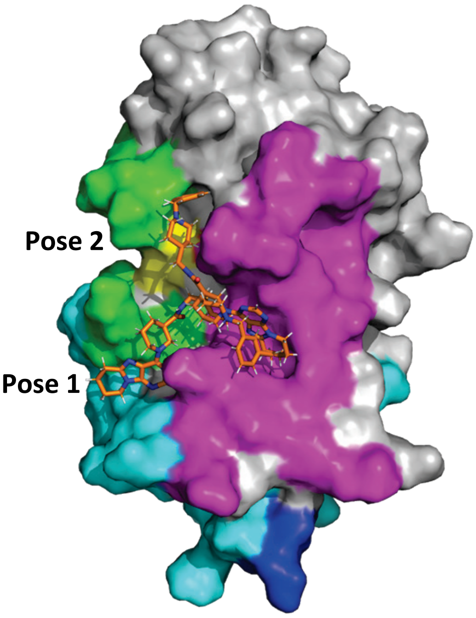


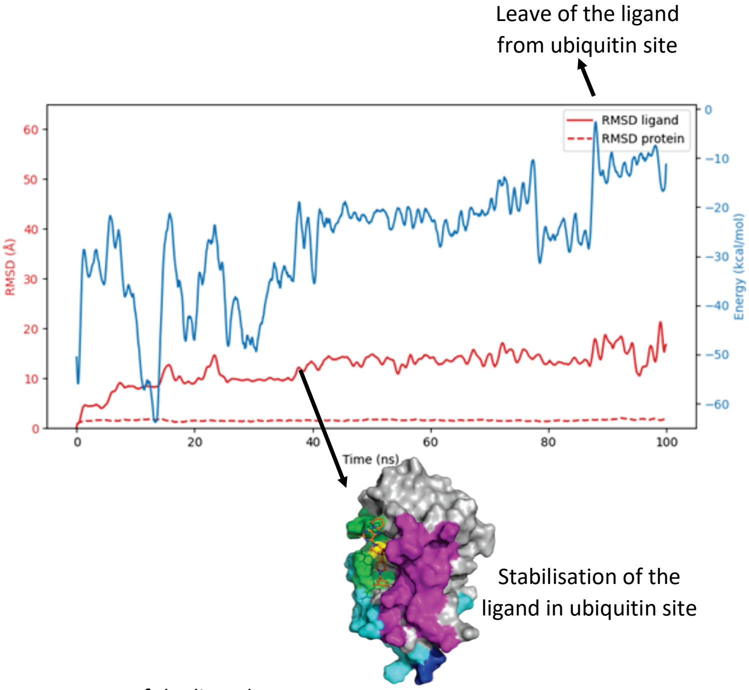


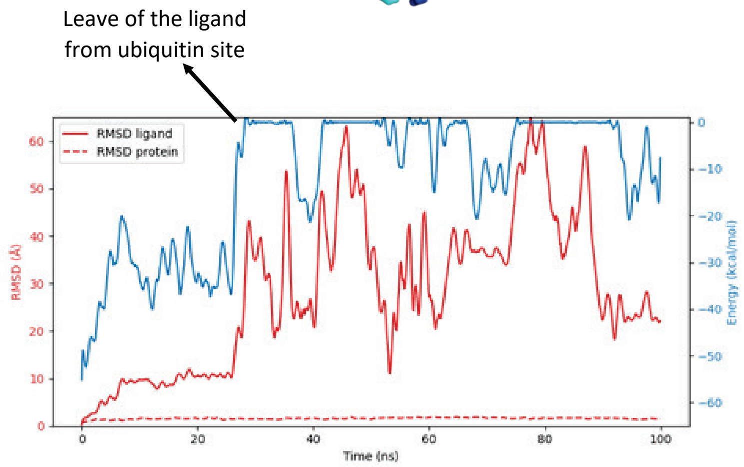


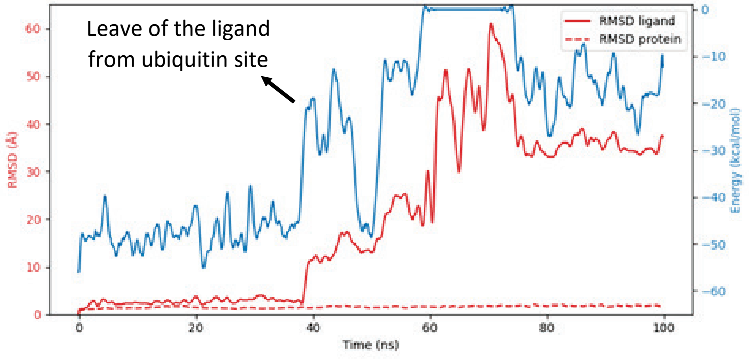

#### ML307: 4ONM ubiquitin and UBE2V2/UBE2V1 site at the same time – pose 2

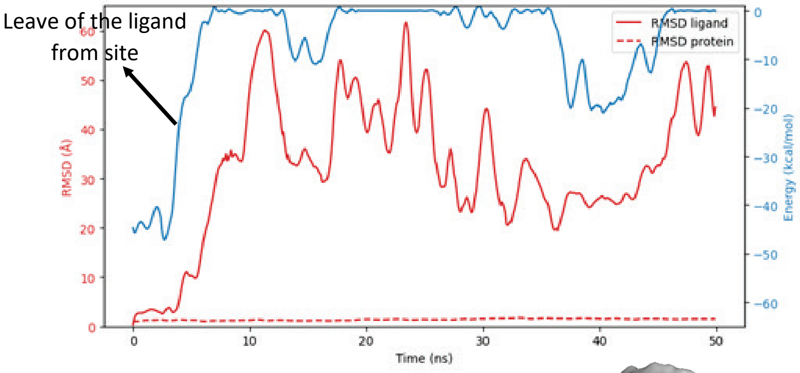


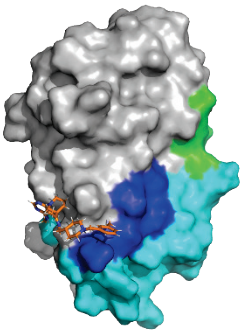

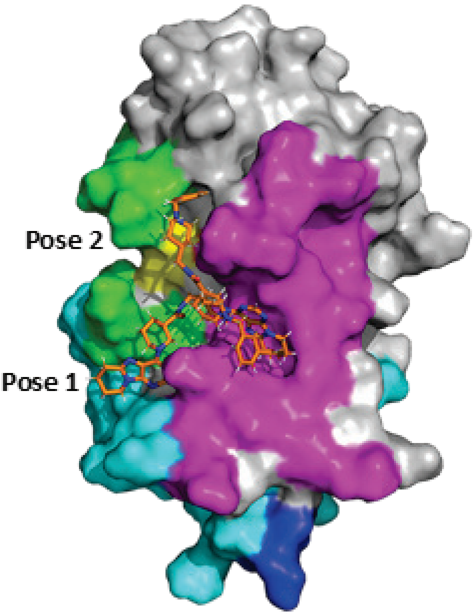


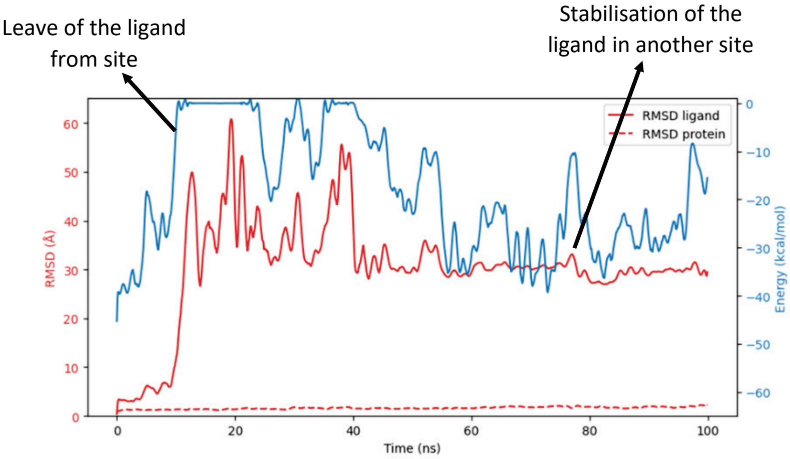


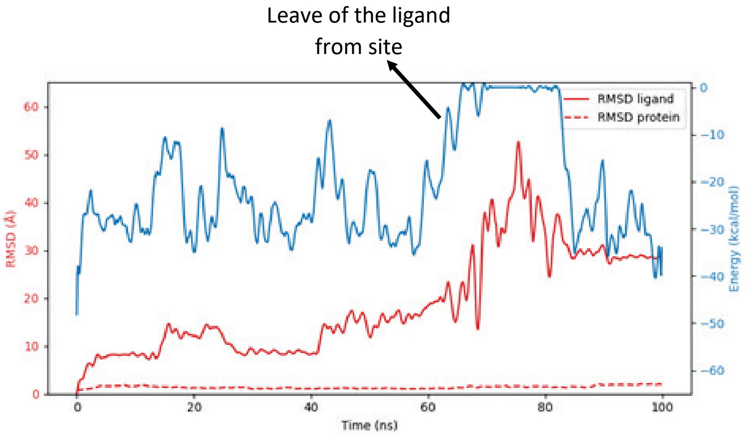

#### ML307: 3HCU – OTUB1 binding site

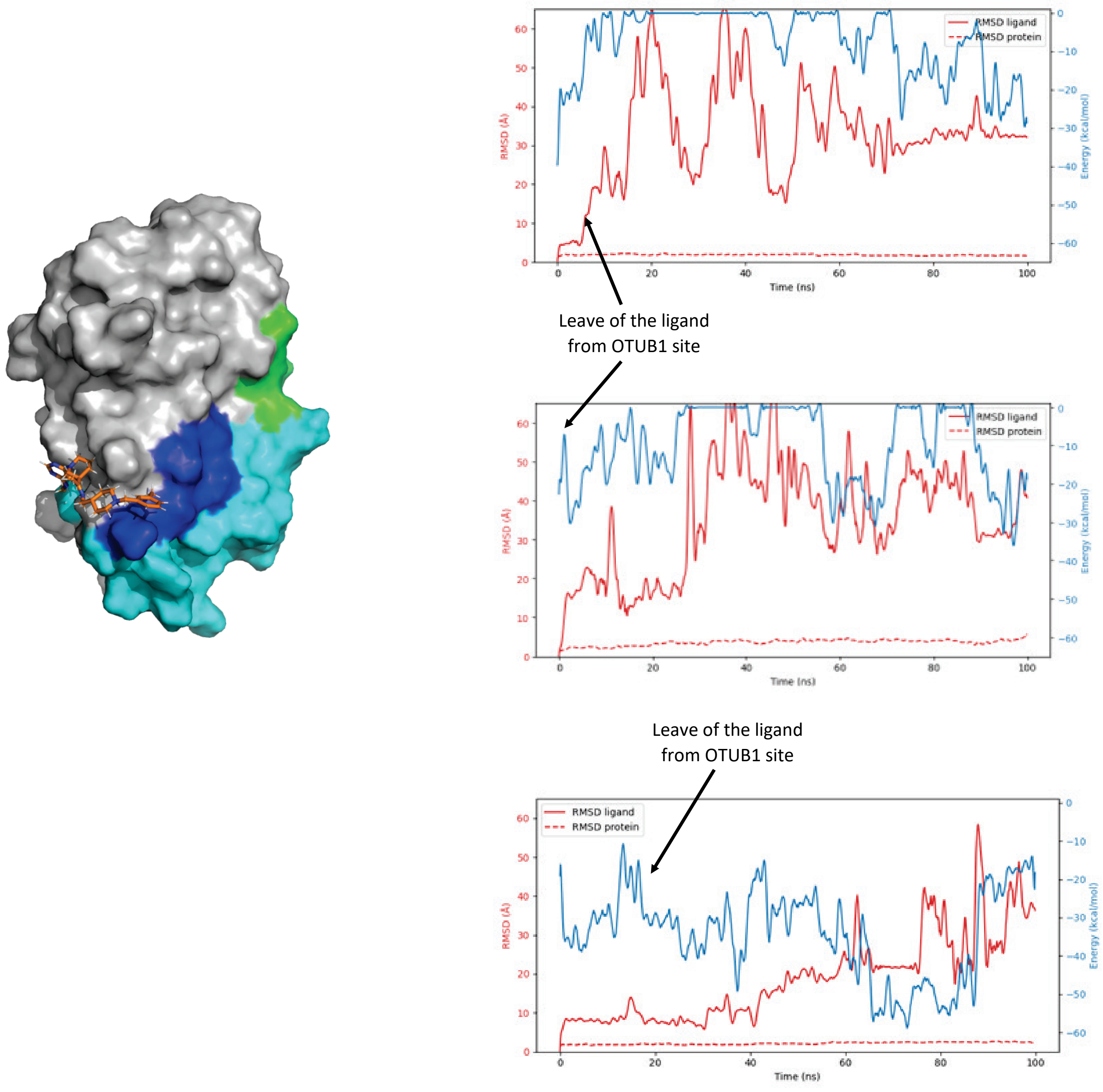


**Figure S3:** The results of the molecular dynamics simulations on the selected poses of ML307 at the ubiquitin site, the UBEV2/UBE2V1 site and OTUB1 binding site.

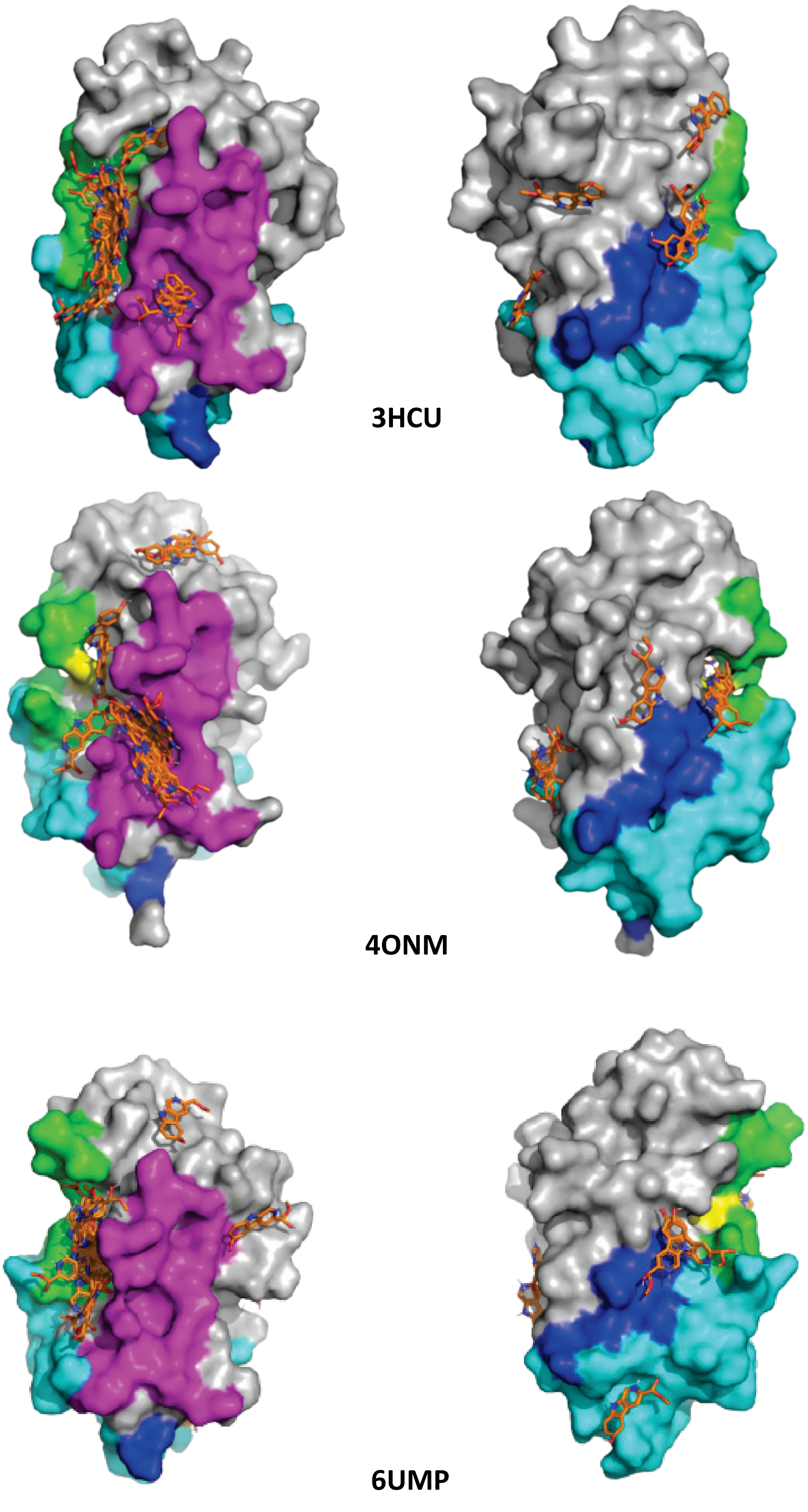


**Figure S4:** The results of the blind docking of Variabine B on the three selected UBE2N X-ray structures selected for the docking study.

#### Variabine B: 6UMP ubiquitin site

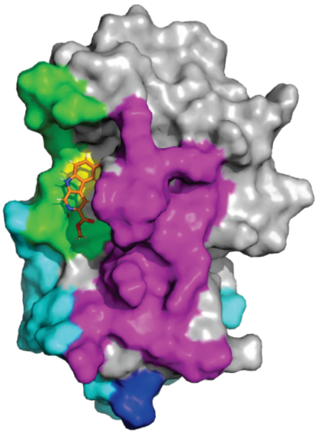

Leave of the ligand from ubiquitin site

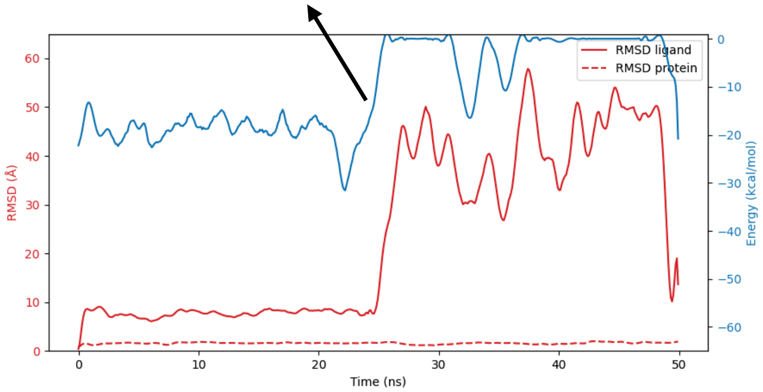


Leave of the ligand from ubiquitin site

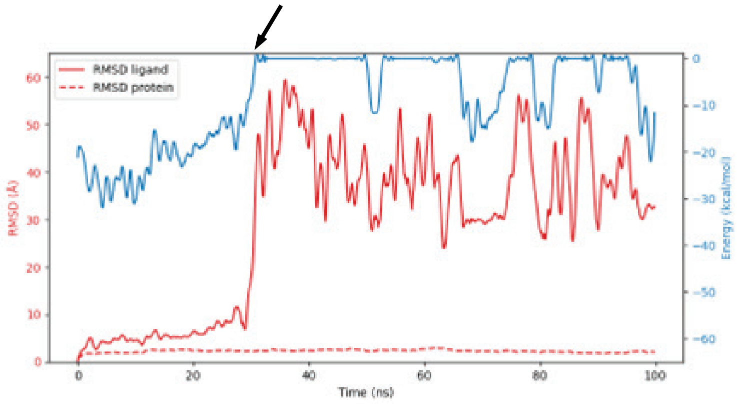


Leave of the ligand from ubiquitin site

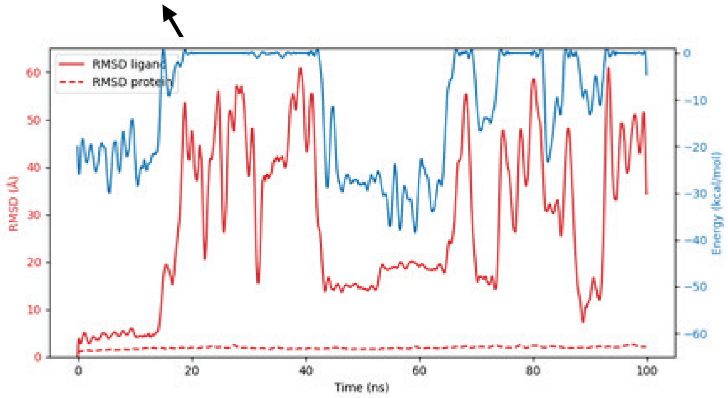

### Variabine B: 4ONM UBE2V2/UBE2V1 site - pose 1

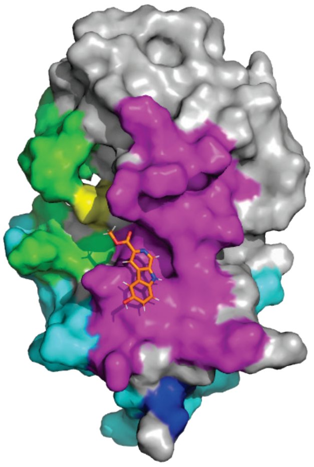

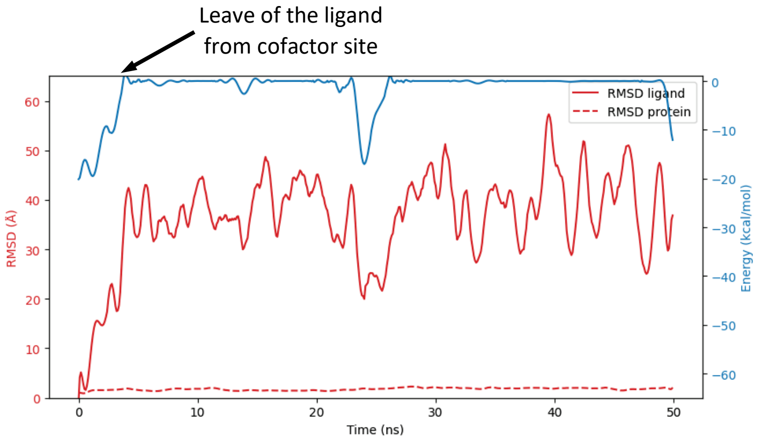


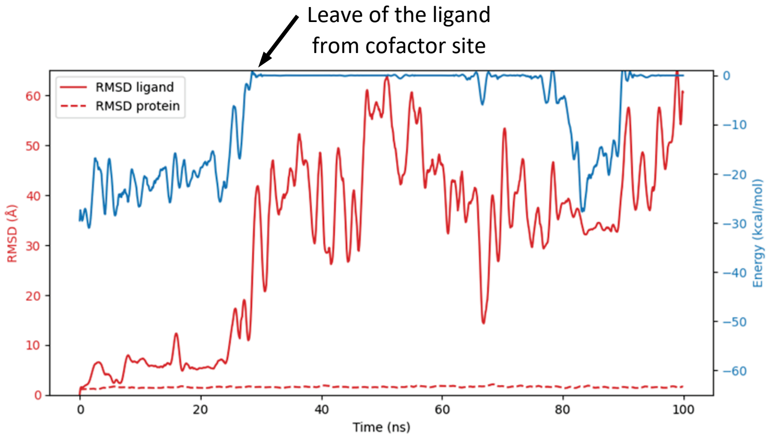


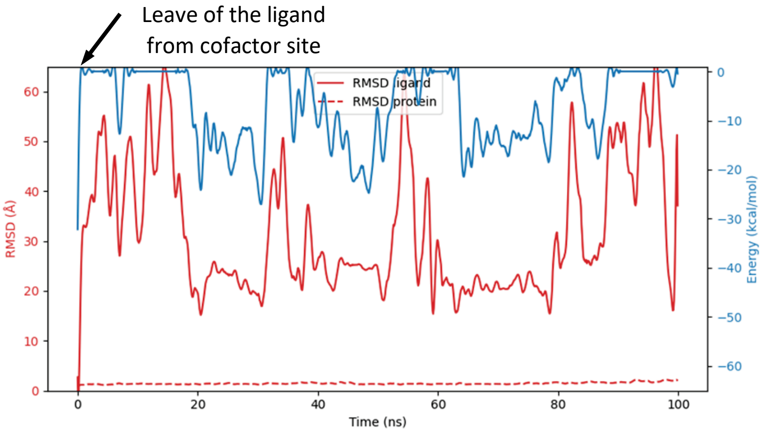

**Variabine B: 4ONM UBE2V2/UBE2V1 site - pose 2**

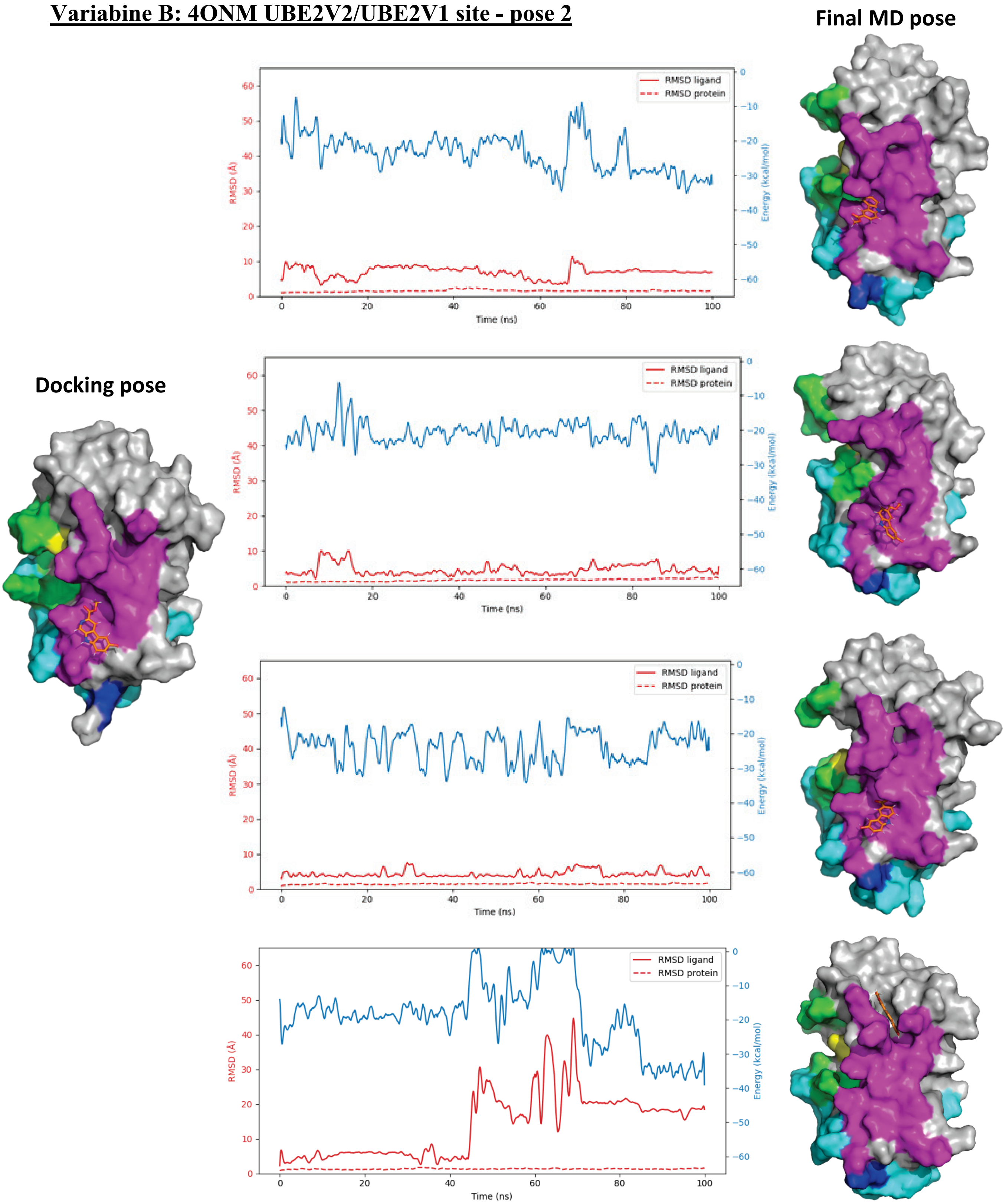


**Figure S5:** The results of the molecular dynamics simulations on the selected poses of Variabine B in the ubiquitin site and in the UBEV2/UBE2V1 site.

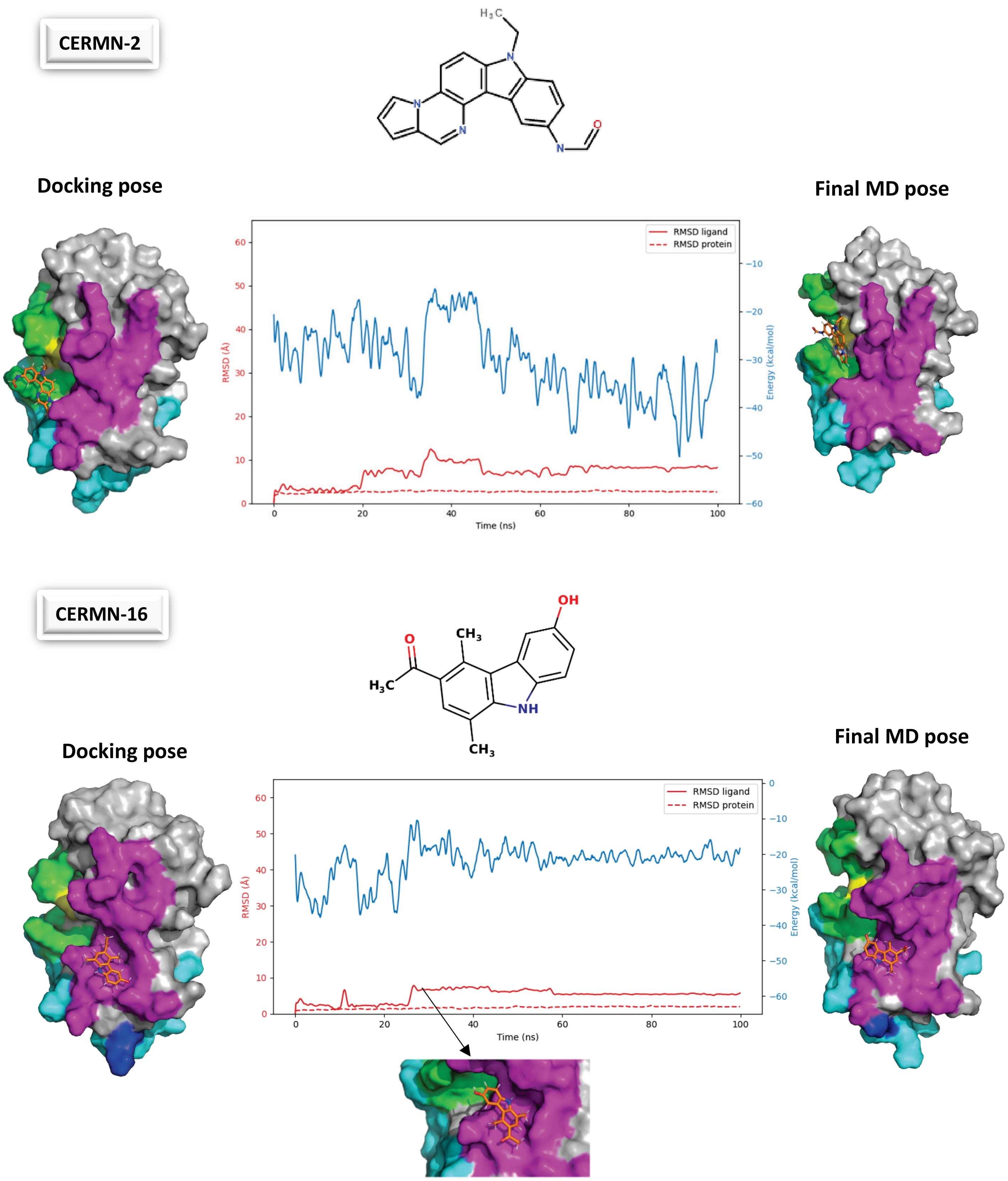


**Figure S6**: Example of the applied molecular screening strategy. (A) CERMN-2 in the ubiquitin-binding site. (B) CERMN-16 in the UBE2V2/UBE2V1 cofactor-binding site.

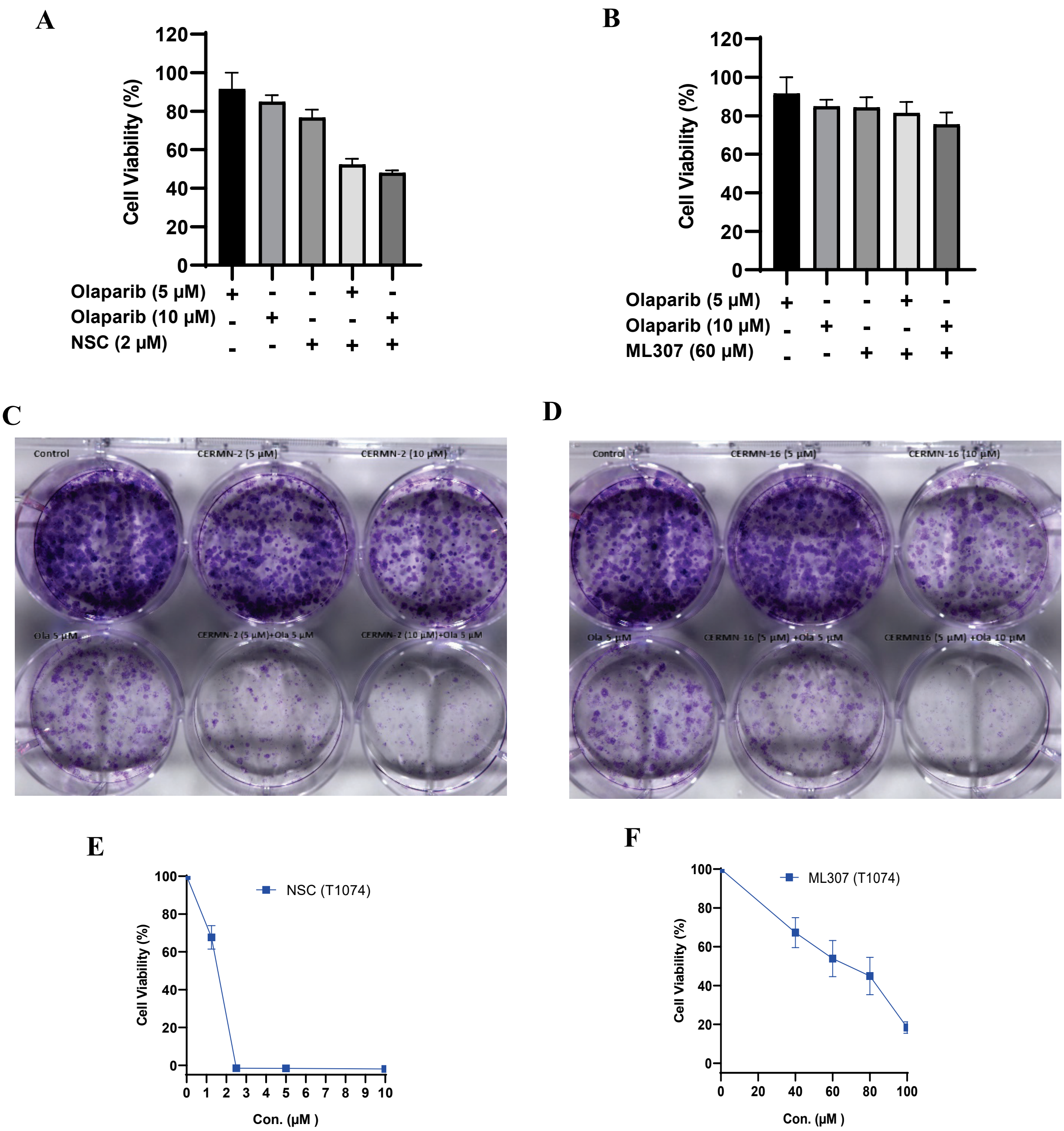


**Figure S7:** (A) Proliferation and sensitization of SKOV3 cells treated with the covalent UBE2N inhibitor NSC697923, the PARP inhibitor Olaparib, and their combination. (B) Proliferation and sensitization of SKOV3 cells treated with the non-covalent UBE2N inhibitor ML307, Olaparib, and their combination. (C–D) Representative images of colony formation in SKOV3 cells treated with the two most potent sensitizing compounds, CERMN-2 and CERMN-16, at concentrations of 5 and 10 µM, alone or in combination with Olaparib (5 µM). The images illustrate the impact of these treatments on cell growth and proliferation; a reduction in colony number and size indicates decreased cell survival and clonogenic potential. (E–F) In vitro cytotoxicity in T1074 epithelial ovarian cells treated in triplicate with NSC697923 and ML307, respectively, at various concentrations.

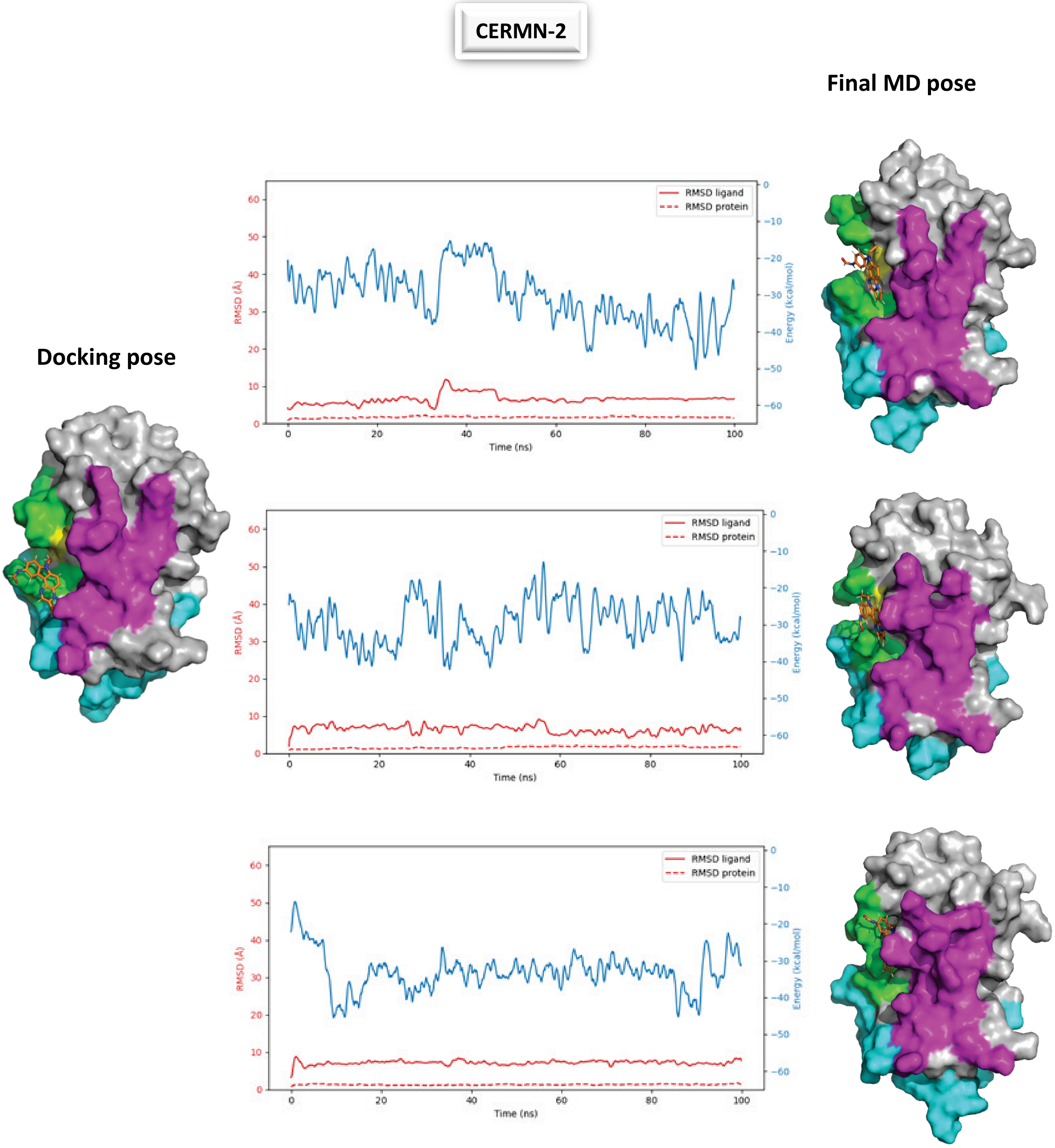


**Figure S8**: The replica of the molecular dynamics simulations on the selected pose of CERMN-2 in the ubiquitin site.

**Synthesis of Variabine B**

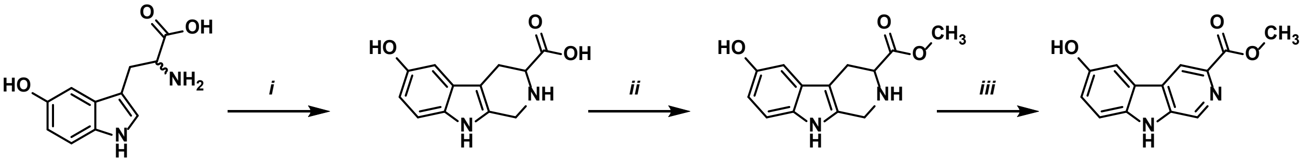

1, 72%    2, 23%    3, 17% Variabine B

**Scheme 1**. Preparation of Variabine B: i. 2-amino-3-(5-hydroxy-1*H*-indol-3-yl)propanoic acid (1 eq.), acetic acid (0.4 eq.), formaldehyde (1.8 eq.), methanol/water, 25°C, 16h; ii. Compound **1** (1 eq.), Hydrochloric acid / MeOH, reflux, 24h; iii. Compound **2** (1 eq.), Pd/C (10 % W/W), xylene, reflux, 4h.

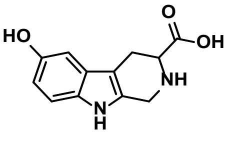

1

6-hydroxy-2,3,4,9-tetrahydro-1*H*-pyrido[3,4-*b*]indole-3-carboxylic acid (**1**) : Following a protocol described by Chiotellis *et al.* [1] (2R/S)-2-amino-3-(5-hydroxy-1*H*-indol-3-yl)propanoic acid;1,3,7-trimethylpurine-2,6-dione (1 equiv., 1g, 4.54 mmol) was dissolved into a mixture of methanol (6 mL) and water (18mL). Then, formaldehyde 37% (1.8 equiv., 607µL, 8.17 mmol) and acetic acid (0.4 equiv., 104µL, 1.82 mmol) were added and the mixture was stirred during 16h at 25°C. The precipitate was filtered and washed with water and ethanol and dried in an oven. The product was obtained as a pale yellow solid and was used without any further purification (760 mg, 72% yield). M.P.: 265-269°C (lit. : 269°C[2]). $^{1}$H NMR (400 MHz, DMSO-$d_6$) $\delta$ : 10.68 (s, 1H), 9.05 (s, 2H), 7.12 (d, $J$ = 8.6 Hz, 1H), 6.78 (d, $J$ = 2.3 Hz, 1H), 6.59 (dd, $J$ = 8.6, 2.3 Hz, 1H), 4.24 (d, $J$ = 15.2 Hz, 1H), 4.16 (d, $J$ = 15.4 Hz, 1H), 3.70 – 3.65 (m, 1H), 3.07 (d, $J$ = 5.0 Hz, 1H), 2.79 (dd, $J$ = 16.1, 10.7 Hz, 1H). 1 proton non-visible under these conditions. $^{13}$C NMR (101 MHz, DMSO) $\delta$ : 169.74, 150.38, 130.63, 127.74, 126.92, 111.54, 111.40, 105.71, 102.01, 56.67, 40.50, 22.86. LC/MS (ESI$^+$): $m/z$ [M + H$^+$] : 233.14, 234.11 [M$^+$ + 2].

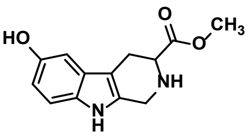

2

[1] Aristeidis Chiotellis et al., "Synthesis, Radiolabeling, and Biological Evaluation of 5-Hydroxy-2-[(18)F]Fluoroalkyl-Tryptophan Analogues as Potential PET Radiotracers for Tumor Imaging," *Journal of Medicinal Chemistry* 59, no. 11 (June 9, 2016): 5324–40, https://doi.org/10.1021/acs.jmedchem.6b00057.

[2] Y. Saiga et al., "Synthesis of 1,2,3,4-Tetrahydro-Beta-Carboline Derivatives as Hepatoprotective Agents. III. Introduction of Substituents onto Methyl 1,2,3,4-Tetrahydro-Beta-Carboline-2-Carbodithioate," *Chemical & Pharmaceutical Bulletin* 35, no. 8 (August 1987): 3284–91, https://doi.org/10.1248/cpb.35.3284.

Methyl 6-hydroxy-2,3,4,9-tetrahydro-1*H*-pyrido[3,4-*b*]indole-3-carboxylate (**2**) : According to a protocol described by Chiotellis *et al.*[1] HCl 1.25 M in methanol was added to a solution of compound **1** (1 equiv., 650 mg, 2.8 mmol) dissolved in methanol (10 mL). The mixture was stirred 24h at reflux. After cooling down, the mixture was evaporated under reduced pressure. The crude was diluted with 20 mL of water and the pH was adjusted to 9 with $NaHCO_3$ and extracted with 3 x 30 mL of EtOAc. The organic layers were dried over $MgSO_4$, filtered and concentrated under reduced pressure. The reaction crude was purified by silica gel column chromatography (DCM/MeOH/$Et_3N$ 98:0:2 to 88:10:2) and concentrated under reduced pressure to afford methyl 6-hydroxy-2,3,4,9-tetrahydro-1*H*-pyrido[3,4-*b*]indole-3-carboxylate (**2**) as a white powder (156 mg, 23% yield). M.P.: 127-129°C (lit. : 130-133°C[3]). $^1$H NMR (400 MHz, DMSO-$d_6$) $\delta$ : 10.35 (s, 1H), 8.52 (s, 1H), 7.03 (d, $J$ = 8.5 Hz, 1H), 6.66 (d, $J$ = 2.3 Hz, 1H), 6.51 (dd, $J$ = 8.5, 2.4 Hz, 1H), 3.94 (d, $J$ = 15.9 Hz, 1H), 3.86 (dt, $J$ = 15.9, 1.9 Hz, 1H), 3.71 – 3.68 (m, 1H), 3.67 (s, 3H), 2.81 (dd, $J$ = 14.8, 4.7 Hz, 1H), 2.70 – 2.61 (m, 1H). 1 proton non-visible under these conditions. $^{13}$C NMR (101 MHz, DMSO) $\delta$ : 173.59, 147.57, 135.25, 130.37, 126.84, 111.52, 110.45, 104.90, 101.38, 56.04, 51.86, 41.65, 25.32. LC/MS (ESI$^+$): *m/z* [M + H$^+$] : 247.21, 248.19 [M$^+$ + 2].

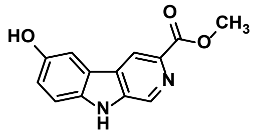


**3**
**Variabine B**

Methyl 6-hydroxy-9*H*-pyrido[3,4-*b*]indole-3-carboxylate – Variabine B (**3**) : According to a protocol described by Cain *et al.*[4] Pd/C (10 % W/W, 90 mg) was added to a solution of **2** (1 equiv., 150 mg, 0.6 mmol) in xylene (15 mL). The mixture was stirred at reflux 16h. After cooling down, the mixture was diluted with methanol (7mL) and filtered on celite pad and washed with a solution of 40 mL of MeOH with 1% $NEt_3$.The solvent was evaporated under reduced pressure and the product was then obtained after recrystallisation of the crude in MeOH. Variabine B was then obtained as an off-white powder (25 mg, 17% yield). M.P.: 258-262°C (lit. : 258-262°C[4]). $^1$H NMR (400 MHz, DMSO-$d_6$) $\delta$ : 11.77 (s, 1H), 9.30 (s, 1H), 8.89 (s, 1H), 8.80 (s, 1H), 7.64 (s, 1H), 7.49 (d, $J$ = 8.8 Hz, 1H), 7.13 (d, $J$ = 8.9 Hz, 1H), 3.90 (s, 3H). $^{13}$C NMR (101 MHz, DMSO) $\delta$ : 166.11, 151.63, 137.92, 135.62, 134.94, 133.65, 127.00, 121.65, 118.78, 117.66, 112.95, 105.99, 51.87. LC/MS (ESI$^+$): *m/z* [M + H$^+$] : 243.15, 244.16 [M$^+$ + 2].

---

[3] Michael Cain et al., .".Beta.-Carbolines: Synthesis and Neurochemical and Pharmacological Actions on Brain Benzodiazepine Receptors," *Journal of Medicinal Chemistry* 25, no. 9 (September 1, 1982): 1081–91, https://doi.org/10.1021/jm00351a015.
[4] Cain et al.
